# Comprehensive multi-platform tyrosine kinase profiling reveals novel actionable FGFR aberrations across pediatric and AYA sarcomas

**DOI:** 10.1101/2023.07.19.548825

**Authors:** Ashleigh M Fordham, Lauren M Brown, Chelsea Mayoh, Alice Salib, Zara A Barger, Marie Wong, Terry C.C. Lim Kam Sian, Jinhan Xie, Kate Gunther, Peter Trebilcock, Rachael L Terry, Paulette Barahona, Pamela Ajuyah, Alexandra Sherstyuk, Anica Avila, Roxanne Cadiz, Callum M Perkins, Andrew J Gifford, Jie Mao, Andrea Zhao, Luke P O’Regan, Daniel Gorgels, Loretta MS Lau, David S Ziegler, Michelle Haber, Vanessa Tyrrell, Richard B Lock, Mark J Cowley, Wayne Nicholls, Roger J Daly, Paul G Ekert, Emmy DG Fleuren

## Abstract

No targeted agents are approved for pediatric sarcomas. Tyrosine kinase (TK) inhibitors represent attractive therapeutic candidates, however, beyond rare TK-activating fusions or mutations, predictive biomarkers are lacking. RNA overexpression of TKs is more commonly observed in pediatric sarcomas, however, an unresolved question is when upregulated TK expression is associated with kinase activation and signaling dependence. We explored the TK molecular landscape of 107 sarcoma patients from the ZERO Childhood Cancer precision medicine program using whole genomic and transcriptomic sequencing. Phosphoproteomic analyses of tyrosine phosphorylation (pY) and functional *in vitro* and *in vivo* assays were also performed in cell lines and patient-derived xenografts (PDXs). Our integrated analysis shows that although novel genomic driver lesions are rare, they are present and therapeutically actionable in selected patients as exemplified by a novel *LSM1-FGFR1* fusion identified in an osteosarcoma patient. We further show that in certain contexts, TK expression data can be used to indicate TK pathway activity and predict TK-inhibitor sensitivity. We exemplify the utility of FGFR-inhibitors in *PAX3-FOXO1* fusion-positive rhabdomyosarcomas (FP-RMS) mediated by high *FGFR4* and *FGF8* RNA expression levels, and overt activation of FGFR4 (FGFR4_pY). We demonstrate marked tumor growth inhibition in all FP-RMS PDXs treated with single agent FGF401 (FGFR4-specific inhibitor) and single agent lenvatinib (multi-kinase FGFR-inhibitor). Clinical benefit of lenvatinib in a relapsed metastatic FP-RMS patient further exemplifies that FGFR-inhibitors deserve additional investigation in FP-RMS patients.

**Statement of significance:** Our multi-omic interrogation of sarcomas in the ZERO Childhood Cancer program illustrates how an RNA-expression biomarker signature (*FGFR4+/FGF8+*) in association with FGFR4 activation identifies that *PAX3-FOXO1*-positive rhabdomyosarcoma patients could benefit from FGFR-inhibitors.

## Introduction

Sarcomas are a heterogeneous group of malignant connective tissue tumors disproportionally affecting the young [1]. Standard-of-care treatment has hardly changed in three decades, consisting of a “one-size-fits-all” approach combining surgery, radiotherapy and (poly)chemotherapy [1–4]. Survival is limited despite aggressive therapy, particularly in the advanced setting (5-20%), and side effects of treatment can be severe [1–4]. No targeted agents are currently approved for pediatric and adolescent and young adult (AYA) sarcoma patients [1], even as more sarcoma genomes are sequenced. This highlights an unmet need for novel and more specific treatments for sarcoma patients and a rationale to look beyond genomic profiles.

One attractive class of cancer targets are the tyrosine kinases (TKs), which are expressed either on the cell membrane (receptor tyrosine kinases (RTKs)) or intracellularly. Aberrant RTK or TK signalling has been implicated in cell autonomous growth signals and repression of apoptosis in pediatric and AYA sarcomas [5–11]. Despite preclinical evidence supporting the utility of certain TK-inhibitors (TKi) for pediatric and AYA sarcoma treatment, clinical trials have not demonstrated efficacy for most patients in the absence of biomarker-driven patient selection [5]. The value of biomarkers is emphasized by precision medicine programs, in which 44-86% of patients receiving a targeted therapy showed sustained responses, including pediatric and AYA sarcoma patients [5, 12]. Although activating genomic TK events are rare in pediatric sarcomas, outlier TK gene expression is much more common and is increasingly recognized as potentially targetable [12, 13]. The clinical utility of gene expression signatures in precision medicine is exemplified by the recent landmark paper by Wong *et al*., where three pediatric sarcoma patients who received a targeted treatment recommendation based on RNA expression levels had sustained responses of up to 50 weeks [12]. A recent example in adult cancer reports efficacy of the FGFR1-4 inhibitor futibatinib in patients with *FGFR*-amplified cancers, not only in patients with structural variants or activating mutations [14]. This suggests that many more pediatric and AYA sarcoma patients might benefit from a TKi based on gene expression levels, distinct from the identification of rare, novel driver lesions using whole genome sequencing (WGS) and transcriptome sequencing (RNAseq). The unresolved question is how to determine when TK gene overexpression truly indicates the potential for therapeutic targeting in the absence of an activating mutation, structural variant or gene amplification.

Our objectives were to investigate the molecular landscape of TKs in pediatric and AYA sarcoma patients from the ZERO Childhood Cancer Precision Medicine Program (ZERO), focusing on previously unrecognized genomic drivers and gene expression profiles [12]. Utilizing WGS and RNAseq data from sarcoma patients, combined with phosphoproteomics and functional *in vitro* and *in vivo* assays in sarcoma cell line models and unique patient-derived xenograft (PDX) models we show that 1) although novel genomic driver lesions are rare, they are present and therapeutically actionable in selected pediatric sarcoma patients, and 2) outlier RNA expression levels of at least one TK are present in the majority of sarcoma patients. Our analysis indicates the utility of a combined *FGFR4+/FGF8+* gene expression signature in association with high FGFR4 activation, detected by phosphoproteomics, to predict sensitivity to novel FGFR-inhibitors (FGFRi) for the treatment of *PAX3-FOXO1*-fusion-positive rhabdomyosarcomas (FP-RMS).

## Methods

Additional details are available in Supplemental Methods. Number of replicates for each experiment are described in the figure legends.

### Patients and samples

The pilot study (TARGET) was approved by the Sydney Children’s Hospitals Network Human Research Ethics Committee (LNR/14/SCH/497) and open from June 2015 to October 2017. The PRISM clinical trial (NCT03336931), conducted as part of ZERO, was approved by the Hunter New England Human Research Ethics Committee of the Hunter New England Local Health District (reference no. 17/02/015/4.06) and the New South Wales Human Research Ethics Committee (reference no. HREC/17/HNE/29). Parents/legal guardian for participants <18 years or participants >18 years of age provided informed consent for study inclusion [12]. Patients with a sarcoma/other soft-tissue malignancy diagnosis were included in this manuscript. Reportable profiles of a subset of these patients were published by Wong *et al.,* 2020 [12] and Lau *et al.,* 2017 and 2021 [15–16].

### WGS data analysis

WGS was conducted at the Kinghorn Centre for Clinical Genomics, Garvan Institute of Medical Research (Australia), using the Illumina HiSeq X Ten platform with a paired-end read length of 150 bases. Methods have been previously detailed, including further downstream processing pipelines for Strelka somatic single nucleotide variants (SNV) and short indel benchmarking, noncoding promoter variation, and variant filtration [12].

### Whole transcriptome data analysis

Whole transcriptome RNAseq was conducted at the Murdoch Children’s Research Institute (Australia) and performed with the TruSeq Stranded mRNA Preparation Kit as previously detailed [12]. Raw gene counts, transcripts per kilobase million (TPM), fragments per kilobase million (FPKM) and isoform expression values were calculated using RSEM (v1.2.31), and fusion detection was called using Arriba (v1.1.0). RNA outlier expression analysis was conducted in R and we note that the reporting of RNA expression has been subject to change over time, as the Z-score is calculated in comparison to the ZERO cohort at time of reporting [17]. All RNAseq expression values are represented as TPM, unless stated otherwise (e.g. Z-score).

### Variant curation

In the ZERO program, variants are classified as ‘likely pathogenic’ or ‘pathogenic’ in accordance with published guidelines [18]. Outlier gene expression levels are defined as Z-score >2 or <−2 as standalone changes, or >1.5 and <−1.5 if supported by expression differences in related genes/pathways. ‘Benign’, ‘likely benign’ or ‘unknown significance’ were not reportable [12].

### Cell culture

All cell lines were cultured according to optimized conditions (Supplemental Table 1), and authenticated by short tandem repeat profiling prior to use.

### *LSM1-FGFR1*, *SPRY2* and *FGF8* overexpression and/or knockout models

*LSM1-FGFR1* and *SPRY2* overexpression models, and *SPRY2* and *FGF8* knockout models were generated as previously described [19]. Supplemental Methods include complete details.

### Western blot

Protein extraction and western blotting was performed as per established techniques [20]. Supplemental Methods contains detailed antibody information.

### Proteome Profiler Arrays

Human Phospho-Receptor Tyrosine Kinase Array Kit (R&D Systems, Cat#ARY001B) and Proteome Profiler Human Phospho-Kinase Array Kit (R&D Systems, Cat#ARY003C) were performed according to manufacturer’s protocols. Signals were visualized on the ChemiDoc Touch System and quantified using Image Lab software v6.1 (Bio-Rad Laboratories).

### Mass Spectrometry

Phosphosproteomic samples were prepared as previously described [21]. Lysates were digested with LysC and trypsin in 4% sodium deoxycholate lysis buffer at pH 7-8. Peptides were cleaned through C18 columns and enriched for pY phosphorylation by pY antibodies (described in Supplemental Methods). Samples were analysed on a Q Exactive HF Mass Spectrometer (Thermo Fisher Scientific) in data-dependent acquisition mode.

### *In vitro* growth inhibition assays

For growth-inhibition assays, cells were seeded into 96-well plates in a final volume of 100µL (sarcoma cell lines) or 200uL (Ba/F3 cells) of normal culture medium. Seeding densities, drug treatments (Selleck Chemicals), incubation times and analysis parameters are described in Supplemental Methods [20].

### IL-3 Withdrawal Assay

GFP positive Ba/F3 cells expressing MSCV, *LSM1-FGFR1* or *FGFR1OP2-FGFR1* (*OP2-FGFR1*) were subjected to an IL-3 withdrawal assay as previously described [19].

### Flow cytometry

Cells were harvested following 8-hour (cell cycle assay) or 96-hour (apoptosis assay) treatment with serial dilutions of inhibitors (Selleck Chemicals) erdafitinib (Cat#S8401), lenvatinib (Cat#S5240) or FGF401 (Cat#S8548). To evaluate cellular apoptosis, samples were stained with Annexin-V and 7-AAD solutions [22]. To investigate cell cycle effects, treated cells were fixed and stained with propidium iodide [22].

### Colony Assays

Cells were seeded on 6-well tissue culture plates. After 48h, serial dilutions of erdafitinib, lenvatinib or FGF401 were added to growth media. Cells were incubated for two weeks (treatments replenished at days 4, 8 and 11) prior to addition of 0.5% crystal violet (Sigma-Aldrich, Cat#C0775) in 50% methanol for 10 min. Crystal violet was removed by repeated rinses with water and plates dried overnight before imaging on the Bio-Rad ChemiDoc Touch System (Bio-Rad Laboratories).

### Establishment of patient-derived xenografts (PDX) or cell line xenografts

Animal procedures were approved under UNSW Animal Care and Ethics Committee (UNSW ACEC 19/82B) and performed using five to nine-week-old female non-obese diabetic/severe combined immunodeficiency/interleukin 2 receptor gamma (null) (NSG) mice [23–24]. For Rh5 cell line xenografts, mice were subcutaneously inoculated with 5×10^6^ tumor cells suspended in 50% matrigel basement membrane matrix (In Vitro Technologies, Cat# FAL356230) in αMEM (Invitrogen, Cat#12571071). For PDX models, 3mm^3^ tumor pieces were subcutaneously implanted [25]. All PDX models were validated by histopathology.

### *In vivo* drug efficacy studies in sarcoma xenografts

Drug treatments were commenced when xenografts reached a volume of 100mm^3^. Tumor growth was measured twice weekly with calipers. Event-free survival (EFS; tumor volume exceeding 400mm^3^) was assessed by Kaplan–Meier analysis with Mantel–Cox testing. Waterfall plots were created to demonstrate maximum decrease in tumor volume from treatment starting volume. Analyses were performed in GraphPad Prism 9.2.0.

### Statistical Analysis

Analyses were performed in GraphPad Prism 9.2.0. One-way ANOVA with Tukey’s multiple comparisons test was performed for all comparisons, except Kaplan–Meier analysis for which Mantel–Cox testing was performed. *: P<0.05, **: P<0.01, ***: P<0.001, ****: P<0.0001. Correlation plots were generated by simple linear regression analysis.

## Data availability

Data supporting the findings of this study are available within the article and supplemental files.

## Results

### Reportable molecular landscape of TK aberrations in pediatric and AYA sarcoma

To establish the frequency of TK activating lesions in high-risk pediatric sarcoma, we analysed WGS and RNAseq data across 58 RTKs and 32 TKs in 107 high-risk pediatric and AYA sarcoma patients enrolled in ZERO, representing over 19 subtypes (“sarcoma cohort,” Table 1). There were 51 patients for whom “reportable” (see Methods) molecular alterations were identified in TKs (Figure 1A, Supplemental Table 2). Although single nucleotide variants (SNVs, three patients, 6%) and structural variants (SVs, five patients, 10%) in TKs of known significance and/or high likelihood of pathogenicity were rare, TK copy number variations (CNVs, twelve patients, 24%) and RNA overexpression (40 patients, 78%) were more commonly reported (Figure 1A). In the majority of pediatric sarcoma patients with reported outlier RNA expression (85%; 34/40), overexpression was not associated with a genomic aberration in the same gene. This includes the elevated *JAK1* and *FGFR4* RNA expression levels characteristic of Ewing sarcoma (EWS) and FP-RMS, respectively (Figure 1A). In only six patients (15%) an associated genomic alteration was identified. This included two fusion-negative rhabdomyosarcoma (FN-RMS; *FGFR4* and *FGFR1*), one osteosarcoma (OS; *PDGFRA*), one malignant peripheral nerve sheath tumor (MPNST; *ALK* and *EGFR*), one angiosarcoma (AS; *VEGFR2*) and one low-grade fibromyxoid sarcoma (FMX; *PDGFRB* and *MET*) patient. RNA overexpression is based on a ZERO cohort outlier analysis (Z-score, calculated on the cohort of all cancer patients enrolled in ZERO) up to the cut-off time for this study.

**Figure 1.**
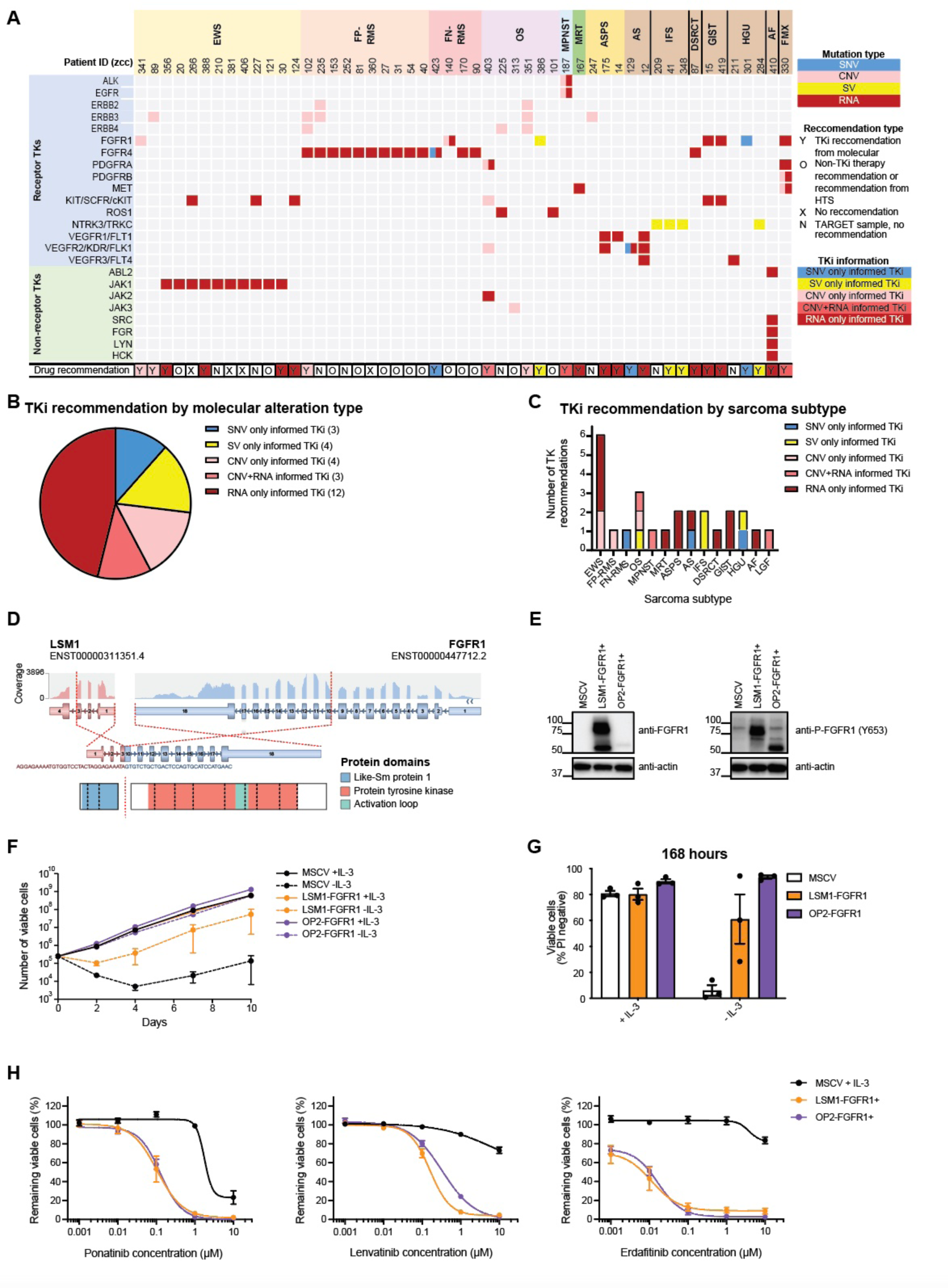
Reportable molecular landscape of TK aberrations in pediatric and AYA sarcoma and novel actionable genomic structural TK variant. A) Reportable molecular TK landscape of sarcoma patients in ZERO. EWS: Ewing sarcoma, FP-RMS: fusion-positive rhabdomyosarcoma, FN-RMS: fusion-negative rhabdomyosarcoma, OS: osteosarcoma, MPNST: malignant peripheral nerve sheath tumor, MRT: malignant rhabdoid tumor, ASPS: alveolar soft part sarcoma, AS: angiosarcoma, IFS: infantile fibrosarcoma, DSRCT: desmoplastic small round blue cell tumor, GIST: gastrointestinal stromal tumor, HGU: high-grade undifferentiated sarcoma, AF: ameloblastic fibrosarcoma, FMX: low-grade fibromyxoid sarcoma. Single nucleotide variants (SNVs) are depicted in blue, copy number variations (CNVs) in pink, structural variants (SVs) in yellow, RNA expression changes in red. In the lower row (drug recommendation), ‘Y’ indicates that a TKi was recommended by the Multidisciplinary Tumor Board (MTB), based on SNVs only (blue), SVs only (yellow), CNVs only (pale pink), CNVs and RNA (dark pink) or RNA only (dark red). ‘O’ indicates a non-TKi therapy MTB recommendation or recommendation from *in vitro* high throughput screening, ‘X’ indicates no MTB therapy recommendation and ‘N’ indicates a TARGET sample, where no MTB recommendations were made. B/C) Summary of the distribution of TKi MTB recommendations based on B) molecular TK profiles (bracketed numbers represent patient numbers) or C) by sarcoma subtype. D) *LSM1-FGFR1* fusion identified in an osteosarcoma patient; (top to bottom) associated transcript and coverage, exon breakpoints, chimeric fusion transcript, sequence around the breakpoint and retained protein domains of the *LSM1-FGFR1* fusion. E) Western blot analysis of *LSM1-FGFR1* and *FGFR1OP2-FGFR1* (*OP2-FGFR1*)-expressing Ba/F3 cells. MSCV (empty vector) control cells were cultured in the presence of IL-3. F) Cell number and viability of *LSM1-FGFR1* and *OP2-FGFR1*-expressing Ba/F3 cells following IL-3 withdrawal over ten-day time course. Total number of viable cells was determined by trypan blue exclusion. Data presented as mean ± standard error of the mean (SEM) (*n=3*). G) Viability analysis of *LSM1-FGFR1* and *OP2-FGFR1*-expressing Ba/F3 cells following 168-hour IL-3 withdrawal. Viability was determined by propidium iodide exclusion, measured by flow cytometry. Data presented as mean ± SEM (*n=3*). H) Resazurin growth-inhibition assay following inhibitor treatment with ponatinib, lenvatinib or erdafitinib in *LSM1-FGFR1* and *OP2-FGFR1*-expressing Ba/F3 cells. Data presented as mean viable cells remining ± SEM (*n=6*).

**Table 1.** Summary of pediatric and AYA sarcoma patient characteristics. Cohort of 107 pediatric and AYA sarcoma patients (“sarcoma cohort”) from the ZERO Childhood Cancer program included in this study. The number of patients, patient age range and gender are summarized for each subtype of sarcoma. The patient diagnosed with “round cell sarcoma” harbors an *EWSR1-PATZ1* fusion as referenced in the text.

| Final Diagnosis | Abbreviation | Number of Patients | Age range | Males | Females |
| --- | --- | --- | --- | --- | --- |
| Alveolar soft part sarcoma | ASPS | 3 | 9-22 | 1 | 2 |
| Angiosarcoma | AS | 2 | 0-6 | 1 | 1 |
| Desmoplastic small round blue cell tumour | DSRCT | 2 | 13-16 | 2 | 0 |
| Ewing's sarcoma | EWS | 29 | 0-20 | 11 | 18 |
| Gastrointestinal neuroectodermal tumour | GNET | 1 | 5 | 0 | 1 |
| Gastrointestinal stromal tumour | GIST | 2 | 10-18 | 1 | 1 |
| High grade undifferentiated sarcoma | HGU | 6 | 2-17 | 5 | 1 |
| Infantile fibrosarcoma | IFS | 3 | 0-2 | 1 | 2 |
| Malignant peripheral nerve sheath tumour | MPNST | 6 | 2-17 | 5 | 1 |
| Malignant rhabdoid tumour | MRT | 8 | 0-3 | 4 | 4 |
| Osteosarcoma | OS | 17 | 9-20 | 10 | 7 |
| Other - Ameloblastic fibrosarcoma | AF | 1 | 18 | 1 | 0 |
| Other - Epithelioid sarcoma | EPI | 1 | 14 | 0 | 1 |
| Other - Leiomyosarcoma | LMS | 1 | 15 | 0 | 1 |
| Other - Low grade fibromyxoid sarcoma | FMX | 1 | 7 | 1 | 0 |
| Other - Round cell sarcoma | ROUND CELL | 1 | 6 | 1 | 0 |
| Other - CIC | CIC | 1 | 14 | 0 | 1 |
| Rhabdomyosarcoma fusion negative | FN-RMS | 7 | 1-18 | 5 | 2 |
| Rhabdomyosarcoma fusion negative, spindle cell | FN-RMS | 1 | 17 | 1 | 0 |
| Rhabdomyosarcoma fusion positive | FP-RMS | 13 | 9-19 | 6 | 7 |
| Synovial sarcoma | SynS | 1 | 7 | 1 | 0 |
| **Total** |  | **107** |  |  |  |

### Treatment recommendations are primarily based on outlier RNA expression

TKi recommendations were rarely based on a TK SNV (*n=3*), SV (*n=4*) or CNV alone (*n=4*) (Figure 1B). These included a heterozygous, activating *VEGFR2* (*KDR*) mutation in the transmembrane domain (c.2312C>G p.Thr771Arg) in an AS patient, an activating *FGFR4* mutation (c.1605C>A p.Asn535Lys) in a FN-RMS patient, and an activating *FGFR1* mutation (c.1638C>A p.Asn546Lys) in a high-grade undifferentiated (HGU) sarcoma (Supplemental Table 2). *NTRK*-associated SVs were identified in four patients which resulted in NTRK-inhibitor recommendations. TKi recommendations were distributed among numerous sarcoma subtypes (Figure 1C).

A novel *LSM1-FGFR1* fusion was identified in an OS patient (Supplemental Table 2). The molecular structure of this fusion suggested it functioned as an oncogenic driver; retaining a coiled-coil domain of LSM1 fused to the TK domain of FGFR1 (Figure 1D, Supplemental Figure 1A). We cloned and expressed *LSM1-FGFR1* in IL-3 dependent Ba/F3 cells. Western blot analysis confirmed expression and activation (marked by FGFR1 phosphorylation) of LSM1-FGFR1, which was sufficient to block apoptosis induced by IL-3 withdrawal, and permit IL-3 independent proliferation (Figure 1E-G, Supplemental Figure 1B). Moreover, Ba/F3 cells dependent on fusion expression died when treated with FGFRi (Figure 1H, Supplemental Figure 1C). This shows that *LSM1-FGFR1* is an oncogenic driver that can be targeted by FGFRi.

TKi therapeutic recommendations most frequently resulted from outlier TK RNA expression, either alone (*n=12)* or in association with CNVs (*n=3)* (Figure 1B). However, fourteen patients for whom an RNA expression change was identified received no related TKi recommendation (Figure 1A, annotation X or O). Currently, the ZERO Multidisciplinary Tumor Board (MTB) rank therapeutic recommendations based on outlier RNA expression alone as the lowest tier ([12] and Supplemental Table 2) because there is little empiric evidence supporting efficacy.

### RNA expression landscape of TKs in pediatric and AYA sarcoma

Given the high prevalence of RNA expression changes in the sarcoma cohort, we aimed to determine when TK RNA expression could be indicative of TK activation, and consequently predictive of TKi response. We analyzed the landscape of TK RNA expression in our cohort to identify expression signatures among sarcoma subtypes independent of reportable profiles and associated MTB therapy recommendations. TK RNA expression data were available for 86 sarcoma patients and two other soft-tissue malignancies (one desmoid tumor and one osteoblastoma, Supplemental Table 3).

Expected associations were identified, including high expression of *JAK1* in EWS (Figure 2A, Supplemental Figure 2A-B), *PDGFR* pathway expression across sarcoma subtypes (Figure 2A, Supplemental Figure 2A) and *FGFR4* expression in FP-RMS (Figure 2A, Supplemental Figure 2A). In some subtypes, links have been established between RNA expression levels and underlying genetic alterations (Figure 1A). Importantly, novel features were also revealed. For example, *PDGFRB* is more frequently overexpressed than *PDGFRA*, with two OS patients expressing the highest levels of *PDGFRB* in the ZERO cohort (Figure 2A, Supplemental Figure 2A,C). A closer examination of other PDGFR-associated genes further revealed upregulation of lesser-known mediators of this pathway, including the *PDGFR-like receptor* (*PDGFRL*) in selected sarcoma patients (Supplemental Figure 2D), and differential expression of PDGF ligands across sarcoma samples, exemplified by high *PDGFC* and *PDGFD* expression in, for example, several OS patients (Supplemental Figure 3A). One gastrointestinal neuroectodermal tumor (GNET, Figure 2A, red arrow) and one FN-RMS (Figure 2A, black arrow) were characterized by outlier levels of *ERBB3*. The GNET patient also expressed high levels of *FYN* (Supplemental Figure 2A). We also observed an overall increase in *PTK7* expression in a variety of sarcoma subtypes compared to other childhood cancers (Figure 2A, Supplemental Figure 3B), most notably in OS, malignant rhabdoid tumor (MRT), MPNST and desmoplastic small round blue cell tumor (DSRCT), and report for the first time outlier expression levels of *PTK7* in a desmoid tumor, with expression levels exceeding any other pediatric cancer profiled in ZERO (Supplemental Figure 2A,3B).

**Figure 2.**
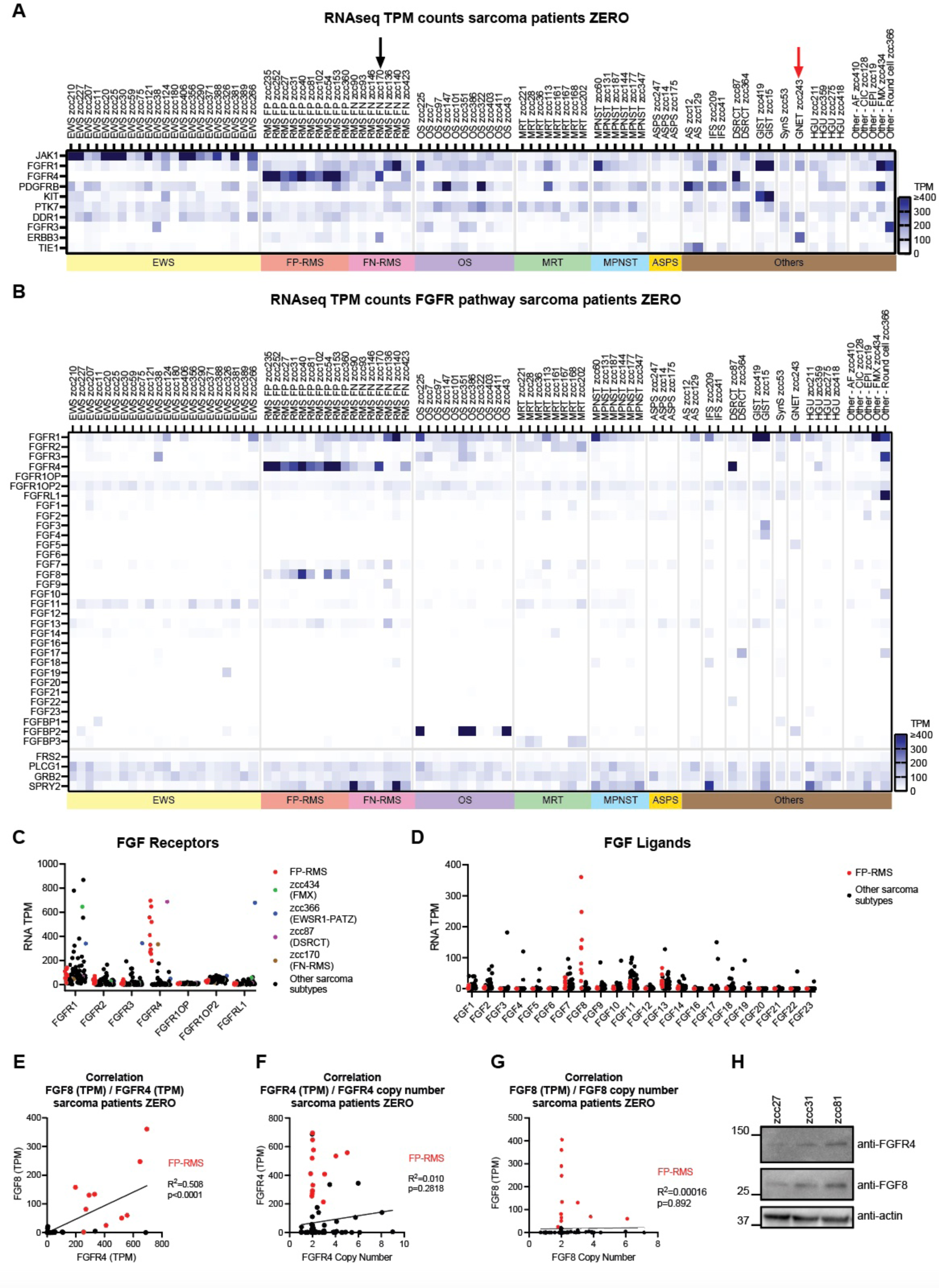
Tyrosine kinase RNA expression in the pediatric and AYA sarcoma cohort: focus on the FGFR-pathway in FP-RMS. A) Heatmap visualizing RNA expression (TPM) with unsupervised hierarchical clustering for TK expression profiles. Top 10 expressed TKs are shown. Black arrow: FN-RMS patient, red arrow: gastrointestinal neuroectodermal tumor (GNET) patient, both characterized by outlier levels of *ERBB3*. B) Heatmap visualizing RNA expression (TPM) of FGFR-pathway genes. C) Dot plot of RNA expression (TPM) for FGF receptors. D) Dot plot of RNA expression (TPM) for FGF ligands. E-G) Correlation between E) *FGFR4* and *FGF8* RNA expression (TPM) F) *FGFR4* (TPM) and *FGFR4* copy number, or G) *FGF8* (TPM) and *FGF8* copy number in ZERO sarcoma patients. Colors indicate specific samples as shown in legends. H) Western blot for FGFR4 and FGF8 in FP-RMS snap frozen PDX tumors. EWS: Ewing sarcoma, FP-RMS: fusion-positive rhabdomyosarcoma, FN-RMS: fusion-negative rhabdomyosarcoma, OS: osteosarcoma, MRT: malignant rhabdoid tumor, MPNST: malignant peripheral nerve sheath tumor, ASPS: alveolar soft part sarcoma, AS: angiosarcoma, IFS: infantile fibrosarcoma, DSRCT: desmoplastic small round blue cell tumor, GIST: gastrointestinal stromal tumor, SynS: synovial sarcoma, GNET: gastrointestinal neuroectodermal tumor, HGU: high-grade undifferentiated sarcoma, AF: ameloblastic fibrosarcoma, CIC: CIC-rearranged sarcoma, EPI: epithelioid sarcoma, FMX: low-grade fibromyxoid sarcoma, Round cell: *EWSR1-PATZ1* round cell sarcoma.

**Figure 3.**
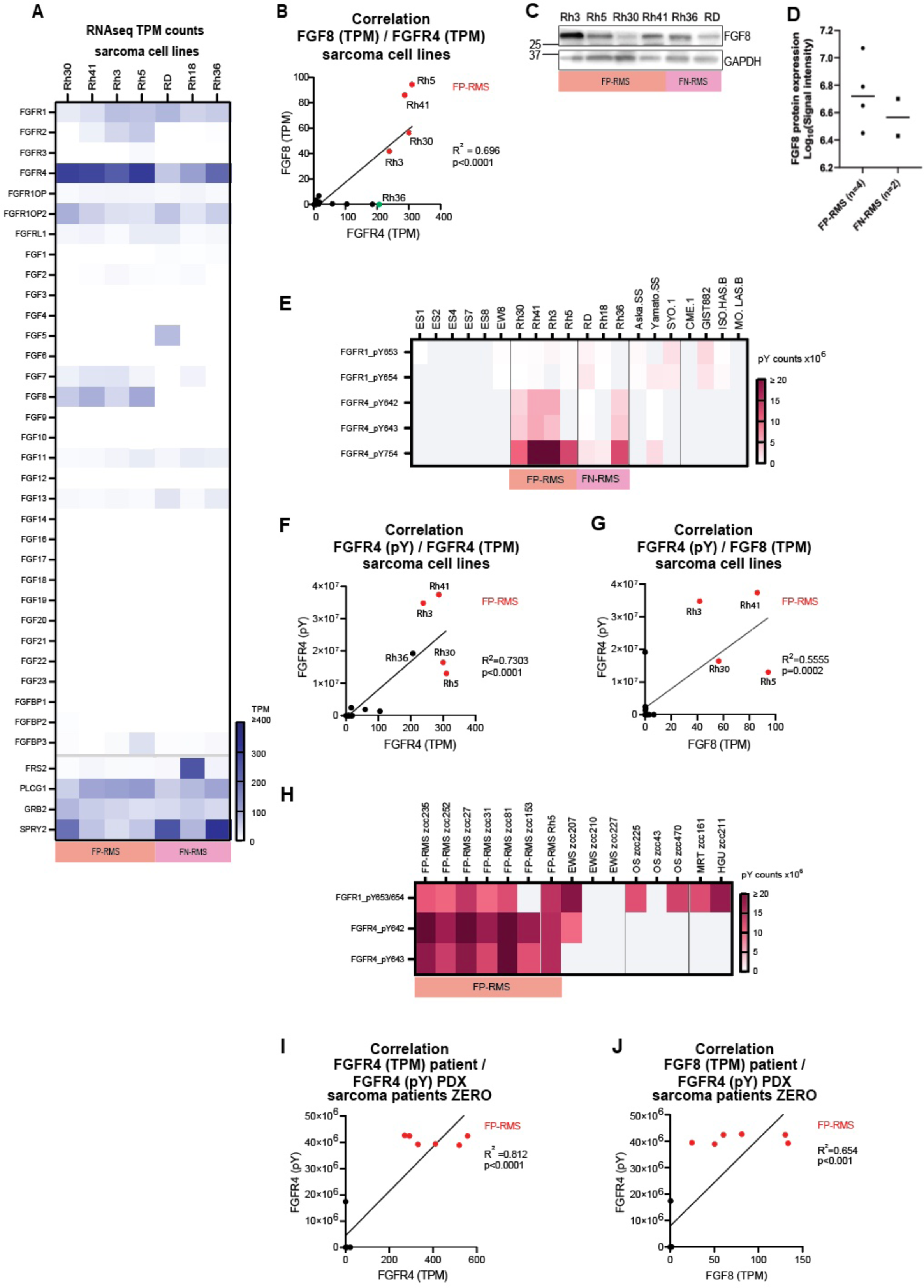
FGFR4 is activated in FP-RMS. A) Heatmap visualizing RNA expression (TPM) of the FGFR-pathway across the RMS cell line panel, ordered by histology and *FGFR* genes. B) Correlation between *FGFR4* and *FGF8* RNA expression across sarcoma cell lines. C) FGF8 Western blot for FP- and FN-RMS cell lines. D) Quantified FGF8 protein expression across RMS cell lines from Western blot shown in C. E) MS quantification of tyrosine residue phosphorylation (pY) of FGFR1 or FGFR4 in sarcoma cell lines. F) Correlation between *FGFR4* RNA TPM and *FGFR4* pY (sum of all pY sites) in sarcoma cell lines. G) Correlation between *FGF8* RNA TPM and FGFR4 pY (sum of all pY sites) in sarcoma cell lines. H) MS quantification of pY of FGFR1 or FGFR4 in sarcoma PDX samples. I) Correlation between *FGFR4* RNA TPM and *FGFR4* pY (sum of all pY sites) in sarcoma PDX samples (*n=13* samples). J) Correlation between *FGF8* RNA TPM and FGFR4 pY (sum of all pY sites) in PDX samples (*n=13* samples). FP-RMS: fusion-positive rhabdomyosarcoma, FN-RMS: fusion-negative rhabdomyosarcoma EWS: Ewing sarcoma, OS: osteosarcoma, MRT: malignant rhabdoid tumor, HGU: high-grade undifferentiated sarcoma.

### The expression of FGFR pathway genes is recurrently altered across pediatric and AYA sarcoma subtypes

Pronounced overexpression of *FGFR*s (Figure 2A, Supplemental Figure 2A) across sarcoma subtypes prompted further analysis of this pathway. We included all FGF ligands, FGF binding proteins (FGFBPs) and other closely related FGF-pathway mediators (Figure 2B-D, Supplemental Table 4). *FGFR1* overexpression was observed across multiple subtypes, including selected cases of FN-RMS, OS, MPNST and gastrointestinal stromal tumor (GIST). For the first time, we report outlier *FGFR1* expression levels in an FMX patient (zcc434) and an *EWSR1-PATZ1* round cell sarcoma patient (zcc366), amongst the highest levels detected in the sarcoma cohort (Figure 2C). The *EWSR1-PATZ1* patient also had the highest levels of *FGFR3* and *FGFRL1* in the sarcoma cohort, suggesting this pathway may be upregulated (Figure 2C). In addition to known association with the FP-RMS subtype (Supplemental Figure 3C), increased *FGFR4* expression was also observed in one DSRCT and one FN-RMS patient (Figure 2C). Another interesting observation was high expression of FGFBP2 in four of the eleven OS patients, top-ranking the sarcoma cohort (Supplemental Figure 3D).

We observed *FGF8* upregulation exclusively in FP-RMS patients. FGF8 is an FGFR4 ligand [26]. FGF8 and FGFR4 have been reported in FP-RMS, although a link between those genes and FGFR-inhibitors had not been thoroughly explored in FP-RMS [27–28]. Overt *FGF8* gene expression was not detected in any other sarcoma or non-sarcomatous childhood cancer patient (Figure 2D, Supplemental Figure 3E). There was a significant positive correlation between *FGFR4* and *FGF8* RNA expression in sarcoma patients, primarily driven by the FP-RMS patients (Figure 2E). Gene expression was not correlated with CNVs for either *FGFR4* (Figure 2F) or *FGF8* (Figure 2G) in the sarcoma cohort, including the FP-RMS. None of the FP-RMS patients with outlier *FGFR4* expression harbored an activating *FGFR4* mutation. Western blot analysis of three FP-RMS patients who had snap-frozen PDX material available confirmed the presence of FGFR4 and FGF8 proteins (Figure 2H). *SPRY2*, a negative regulator of the FGFR-pathway, was highly expressed in two FN-RMS patient samples, and to a lesser extent also in one IFS and one HGU patient (Figure 2B).

### *FGFR4* and *FGF8* RNA expression levels correlate with FGFR4 protein activation status in RMS

Wildtype (WT) *FGFR4* overexpression has been reported in FP-RMS before. However, *in vivo* targeting of WT *FGFR4* in FP-RMS models with drugs like ponatinib yielded disappointing results, leading to the consensus that targeting WT *FGFR4* in FP-RMS is futile [29]. We hypothesized that having *FGF8* concurrently overexpressed with *FGFR4* in FP-RMS, together with FGFR4 activation, might predict FGFRi-sensitivity in FP-RMS.

We set out to test these hypotheses in a panel of sarcoma cell lines, including FP- and FN-RMS, which demonstrated similar *FGFR*-pathway RNA expression profiles to patient samples (Figure 3A, Supplemental Figure 4A, Supplemental Table 5) including high *FGFR4* in all four FP-RMS cell lines and one FN-RMS cell line, and high *FGF8* expression exclusive to the four FP-RMS cell lines. The positive correlation between *FGFR4* and *FGF8* RNA expression detected in patient samples was also observed in sarcoma cell lines, where again the FP-RMS cells were driving this correlation (Figure 3B). FGF8 protein was present in FP- and FN-RMS cell lines, though overall, FGF8 protein expression was lower in FN-RMS versus FP-RMS (Figure 3C-D). pY Mass-Spectrometry analysis (MS) showed FGFR4 tyrosine phosphorylation (FGFR4_pY) in all four FP-RMS cell lines and in the FN-RMS cell line Rh36 (Figure 3E) [10]. There was a positive correlation between FGFR4_pY and both *FGFR4* (Figure 3F) and *FGF8* (Figure 3G) RNA expression in sarcoma cell lines. At the protein level, FGF8 positively correlated with FGFR4_pY (Supplemental Figure 4B). We also show evidence of FGFR4 activation in all tested FP-RMS PDX models established from ZERO patients (Figure 3H) and a positive correlation of FGFR4_pY with *FGFR4* (Figure 3I) and *FGF8* (Figure 3J) RNA expression. PDX models of EWS, OS, MRT and HGU examined concurrently had low or no detectable FGFR4_pY (Figure 3H), and the corresponding patient RNA expression was low for *FGFR4* and *FGF8* (Figure 2B). Together, these data indicate that both *FGFR4* and *FGF8* are closely associated with FGFR4 activation.

### Novel FGFRi have potent effects on FP-RMS cell viability at clinically relevant doses

To determine whether FP-RMS cells with the *FGFR4+/FGF8+* gene expression signature, in association with FGFR4_pY activation are more sensitive to FGFRi than cells which lack this expression pattern, we exposed FP-RMS cells lines to lenvatinib (VEGFRs, PDGFRs, FGFRs), erdafitinib (FGFR1-4), AZD4547 (FGFR1-4) and FGF401 (FGFR4 only). Ponatinib (pan-kinase, targets include FGFRs) was included as a reference [29] (Figure 4, Supplemental Figure 5A). As an FGFRi-sensitive positive control, we included a pre-B cell line (Ba/F3) engineered to express the *FGFR1OP2-FGFR1* (*OP2-FGFR1*) driver [30]. Additional cell lines from other sarcoma subtypes, including two FN-RMS cell lines, were included as (negative) controls. As expected, none of the cell lines were as sensitive to FGFR-targeted agents as the positive control (OP2-FGFR1). Three out of the four FP-RMS cell lines (Rh3, Rh5 and Rh41) were more sensitive to all inhibitors than the FN-RMS or any of the other sarcoma cell lines (Figure 4A). This includes the FN-RMS Rh36 and OS MG63 cell lines, that exhibit *FGFR4* RNA expression levels similar to the FP-RMS cell lines, but lack *FGF8* expression (Figure 3A, Supplemental Figure 4A). This suggests that FGF8 could be important in mediating FGFRi- sensitivity in FP-RMS. The FGFR4-specific inhibitor FGF401 killed more FP-RMS cells than FN-RMS cells, indicating that the activity of the less-specific FGFRi are likely also mediated through FGFR4. Importantly, FGF401 did not affect the OP2-FGFR1 cells (due to lack of activity against FGFR1). Apart from ponatinib, the effects on cell viability were observed at concentrations within clinically achievable ranges (defined as below the maximum single dose (Cmax) and steady state (Css) concentrations achievable for a patient for each drug) (Figure 4A). When drug efficacy was assessed by analyzing area under the curve (AUC), we observed a significant difference between the three responding FP-RMS cell lines and other sarcoma cell lines (Figure 4B).

**Figure 4.**
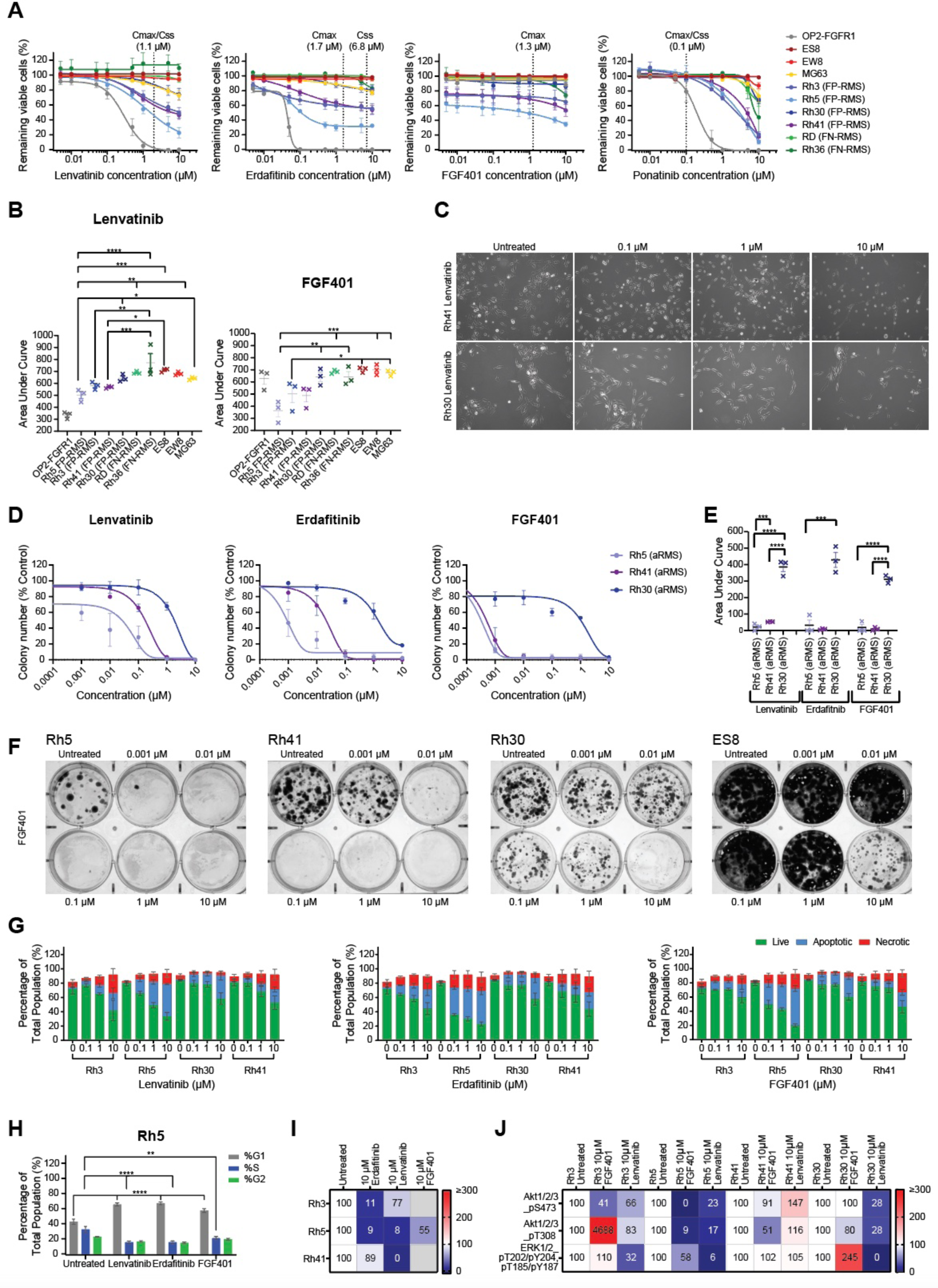
FP-RMS cell lines are sensitive to FGFRi *in vitro*. A) Resazurin growth-inhibition assay following 76-hour inhibitor treatment with lenvatinib, erdafitinib, FGF401 or ponatinib in sarcoma cell lines. An *OP2-FGFR1* overexpression model was included as a positive control for non-FGFR4-specific inhibitors. Data presented as mean percentage of viable cells remaining ± SEM (*n=3*). B) Area under the curve (AUC) was calculated (from A), lenvatinib and FGF401 assays shown. *: P < 0.05, **: P < 0.01, ***: P < 0.001, ****: P < 0.0001 (one-way ANOVA with Tukey multiple-comparisons test). Data presented as mean AUC ± SEM (*n=3*). C) Representative images captured (*n=3* from A) 76- hours after treatment, Rh41 and Rh30 lenvatinib-treated cells shown. D) Colony assay following two-weeks of treatment with lenvatinib, erdafitinib or FGF401 in Rh5, Rh41 and Rh30 FP-RMS cells (mean colony number ± SEM, *n=3*). E) AUC for colony assay (from D) was calculated. *: P < 0.05, **: P < 0.01, ***: P < 0.001, ****: P < 0.0001 (one-way ANOVA with Tukey multiple-comparisons test). Data presented as mean AUC ± SEM (*n=3*). F) Representative images of colony assay (*n=3* from D-E), FGF401-treated colonies shown. G) Flow cytometry-based Annexin-V/7-AAD analysis. FP-RMS cell lines cells were harvested following 96-hour treatment with lenvatinib, erdafitinib or FGF401 and stained with Annexin-V and 7-AAD. Gating was performed against positive controls for live (Annexin-V and 7-AAD negative), apoptotic (Annexin-V positive) or necrotic (7-AAD positive and Annexin-V negative) cells (gating strategy example: Supplemental Figure 5D) and applied to cells which were untreated or treated with FGFR4 inhibitors. Data presented as mean percentage of live, apoptotic or necrotic cells present in the total population ± SEM (*n=3*). H) Cell cycle analysis of Rh5 cells treated for 8-hours with 10 µM lenvatinib, erdafitinib or FGF401, data presented as mean percentage of G1, S and G2 phase cells present in the total population ± SEM (*n=3*). **: P < 0.01, ***: P < 0.001, ****: P < 0.0001 when compared to untreated control (one-way ANOVA with Tukey multiple-comparisons test). I) RTK and J) phospho-kinase arrays demonstrating relative change in protein phosphorylation of FGFR4 or downstream AKT and ERK pathway kinases upon treatment for 8-hours with 10 µM lenvatinib, erdafitinib or FGF401. Phosphorylation was normalized to control signals on the membrane, and to arrays performed on untreated controls.

We noted that the colorimetric viability assays may underestimate the true effects of the drugs on sensitive FP-RMS cells (Supplemental Figures 6-9) For example, the microscopic appearance of Rh41 FP-RMS cells suggested that many cells were dead or dying after lenvatinib treatment at 1 and 10 µM (Figure 4C). To pursue this further, we performed long-term colony assays on three FP-RMS lines of varying susceptibility to lenvatinib, erdafitinib and FGF401, to measure colony forming potential in the presence of FGFRi. These data confirmed that Rh5 and Rh41 cells were more sensitive to FGFRi than indicated by the short-term viability assays (Figure 4D-E), most noticeable with the FGFR4-specific inhibitor FGF401 (Figure 4F) which markedly reduced colony formation even at 0.01 µM in both Rh5 and Rh41, while Rh30 cells were significantly more resistant (p<0.001).

To explore whether lenvatinib, erdafitinib and FGF401 were inducing apoptotic cell death in FP-RMS, we exposed each of the four cell lines to FGFRi for 96-hours and performed a flow cytometry-based Annexin-V/7-AAD analysis (Figure 4G). These data showed an expected decline in the live cell population after 96-hours. Treated cells showed increased Annexin-V expression compared to untreated controls, indicating apoptotic cell death. Inhibition of oncogenic signaling may also induce cell cycle arrest prior to cell death. To assess this, we exposed each of these cell lines to 10 µM FGFRi for 8-hours and analyzed the proportion of cells in G1, S and G2/M phase by flow cytometry. These data show that all three agents induced G1 cell cycle arrest (p<0.0001, Figure 4H) prior to apoptosis (Figure 4G). Using antibody arrays, we further showed that each FGFRi showed on-target effects by reducing FGFR4_pY (Figure 4I). Downstream phosphorylation events on AKT1/2/3 and ERK1/2 were affected to various degrees by the different drugs. Overall, the most pronounced decreases in FGFR4_pY and downstream mediators with all drugs were observed in Rh5, the best responding cell line (Figure 4J).

### FGFRi sensitivity is mediated via FGF8

Increased susceptibility to FGFRi was observed exclusively in FP-RMS, which are characterized by *FGFR4* and *FGF8* co-expression. The lack of FGFRi efficacy in sarcoma cell lines expressing *FGFR4* but not *FGF8* suggests that *FGF8* is a biomarker of response. To functionally test the importance of FGF8 in FP-RMS cells, we examined the effects of exogenous FGF8 ligand addition to Rh5, Rh41 and Rh30 FP-RMS cell cultures, as well as by examining the effects of *FGF8* CRISPR-Cas9 knock-out (KO) in Rh5 FP-RMS cells. A two-week exposure to FGF8 increased proliferation in two out of three FP-RMS cells (Supplemental Figure 10A). Subsequently, drug response assays were performed in naïve or FGF8 pre-conditioned Rh5 and Rh30 cells. In short-term viability assays, FGF8 pre- conditioning did not significantly alter the response to FGF401, lenvatinib or erdafitinib (Figure 5A), however long-term colony assays indicated that FGF8 pre-conditioning did result in a greater decrease in colony forming capacity following erdafitinib treatment (Figure 5B-C). *FGF8* KO dramatically affected proliferation of Rh5 cells in both short- and long-term assays (Supplemental Figure 10B-D), supporting a critical role for FGF8 in FP-RMS progression and precluding reliable drug sensitivity testing in this model.

**Figure 5.**
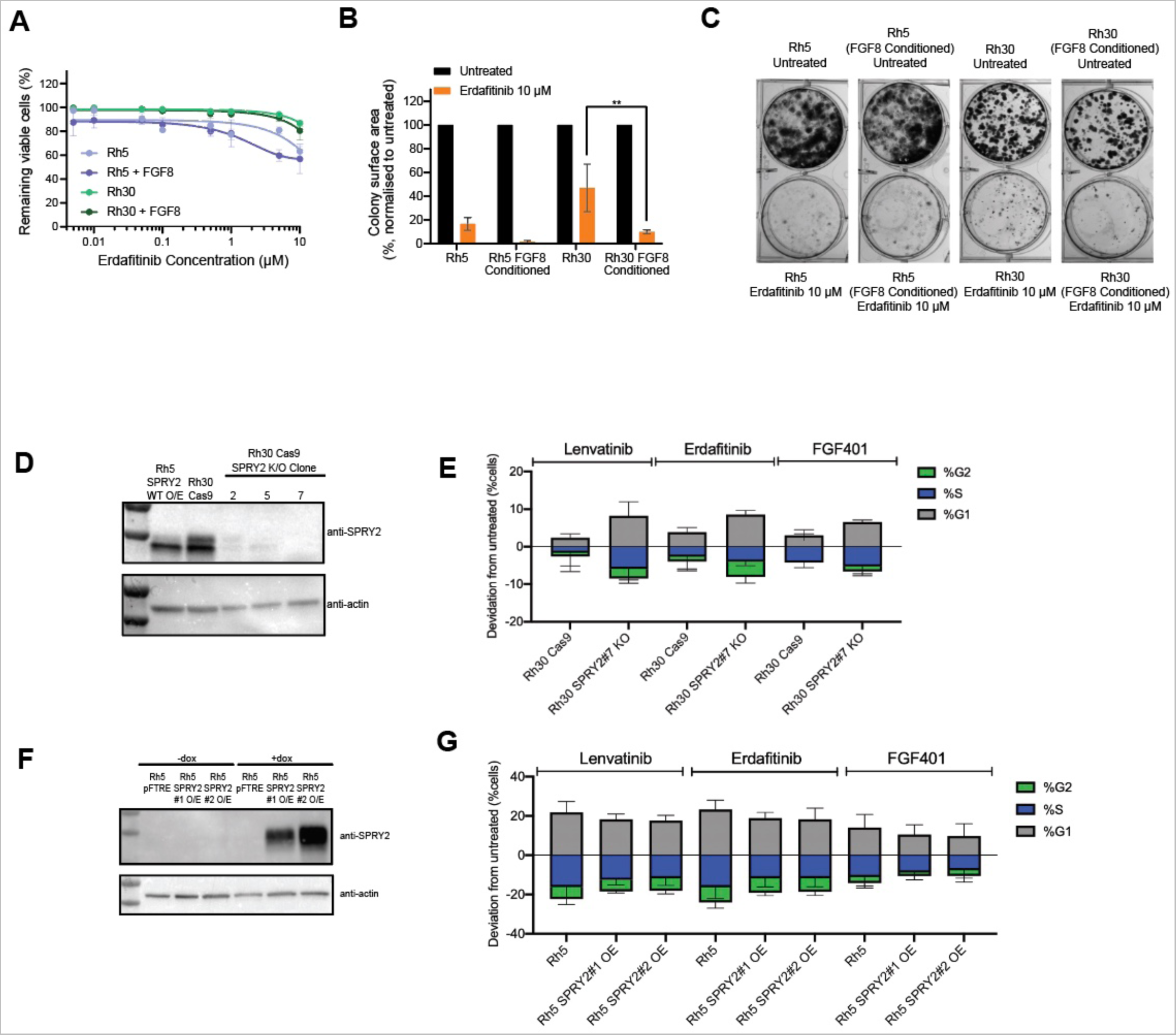
FGF8 is a mediator of sensitivity to FGFRi in FP-RMS. A) Resazurin growth-inhibition assays after two-weeks of FGF8 pre-conditioning, performed after 76-hour treatment with erdafitinib, in the presence or absence of 20 ng/mL FGF8. Data presented as mean viable cells remaining ± SEM (*n=3*). B) Colony assay following two-weeks 10 µM erdafitinib treatment. Data presented as mean colony number ± SEM (*n=3*). **: P < 0.01 when compared to untreated control (one-way ANOVA with Tukey multiple-comparisons test). C) Representative images of colony assay (*n=3* from B). D) Rh30 *SPRY2* KO models were generated using CRISPR-Cas9, as previously described. Bulk cell populations were single cell sorted to select clonal populations (2, 5, 7) with complete knockout, which was confirmed by Western blot analysis. Complete KO was observed in clone 7 and as such this was used for further functional studies. Rh5 cells with enforced overexpression of SPRY2 were used as a positive control (Rh5 SPRY2 #1 O/E). E) Cell cycle analysis of Rh30 SPRY2 knockout clone 7 and Cas-9 control treated for 8-hours with 10 µM lenvatinib, erdafitinib or FGF401. Data presented as the percentage of drug-treated cells in G1, S and G2 phase normalized to untreated control of each respective cell line construct ± SEM (*n=3*). F) Rh5 SPRY2 overexpression (OE) models were generated and protein overexpression confirmed by Western blot. Models include SPRY2 wildtype (WT; SPRY2#1) OE, and the SPRY2 P106S variant (MUT; SPRY2#2) OE, the latter representing a SPRY2 variant observed in the normal population and predicted to have no functional effect. G) Cell cycle analysis of two SPRY2 overexpression Rh5 models (SPRY2#1 OE, SPRY2#2 OE) and pFTRE control treated for 8-hours with 10 µM lenvatinib, erdafitinib or FGF401. Data presented as percentage of drug-treated cells in G1, S and G2 phase normalized to untreated control of each respective cell line construct ± SEM (*n=3*)

We also observed that Rh30 cells, the only FP-RMS cell line that did not respond to any of the FGFRi (Figure 4A-F), expressed high levels of the negative pathway mediator *SPRY2* (Figure 3A). We hence also functionally addressed the role of SPRY2 in FP-RMS. We do however note that in our clinical cohort, none of the included FP-RMS patients exhibited high *SPRY2* levels (Figure 2B). *SPRY2* expression and its potential role in conferring FGFRi-resistance is as such considered to be of much less clinical relevance compared to our findings on *FGF8,* which is a recurring molecular feature in FP-RMS patients. We generated *SPRY2* CRISPR- Cas9 KO Rh30 cells, and SPRY2-overexpressed (OE) Rh5 cells (Figure 5D and F). Neither SPRY2 KO nor OE altered FGFRi sensitivity in our tested models in short-term viability assays (Supplemental Figure 10E-F). However, cell cycle analysis did show a trend towards increased sensitivity to FGFR-targeting drugs in Rh30 *SPRY2* KO cells versus their isogenic control, exemplified by increases in G1 arrest and decreases in S-phase and G2 (Figure 5E). We observed the opposite trend in the Rh5 SPRY2 OE models versus their isogenic control (Figure 5G). This suggests that although SPRY2 may not be a critical determinant on its own in mediating FGFRi sensitivity or resistance, it appears to be partly involved.

Collectively, these data are consistent with our hypothesis that the FP-RMS subtype is susceptible to FGFRi, which is largely mediated by FGF8.

### Lenvatinib and FGF401 are effective in FP-RMS *in vivo* models

We next tested lenvatinib, erdafitinib, FGF401 and ponatinib as single agents in Rh5 xenografts *in vivo*. While ponatinib and erdafitinib had limited effect on tumor growth, there was marked single-agent efficacy of both FGF401 (FGFR4-specific inhibitor) and lenvatinib (VEGFR-, PDGFR-, FGFR-inhibitor), with inhibition of tumor growth observed in three out of four mice throughout the treatment period, significantly extending event-free survival (EFS) (Figure 6A- C).

**Figure 6.**
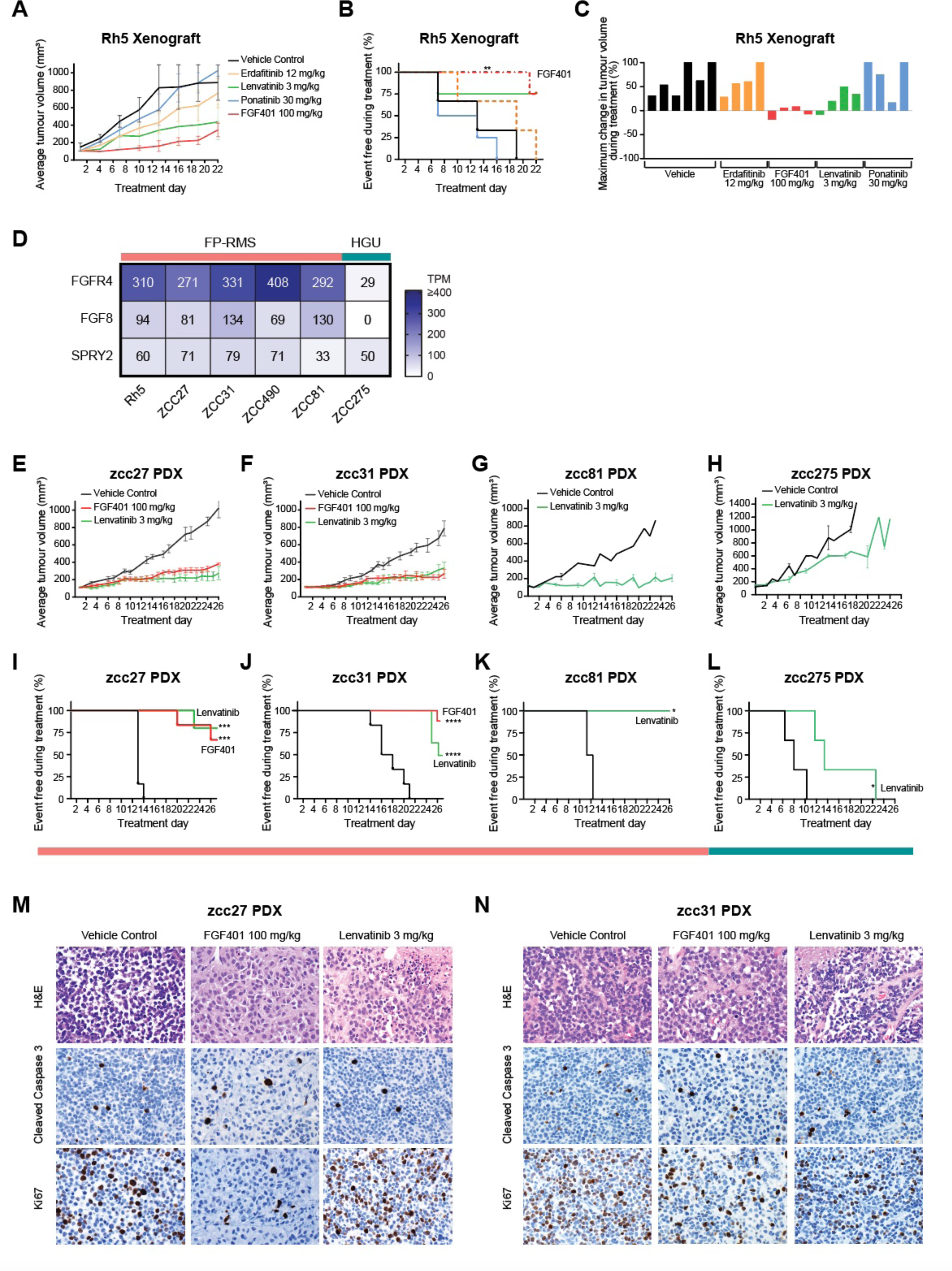
FP-RMS cell line and PDX models are responsive to FGFRi *in vivo*. A) Tumor growth (mean tumor volumes ± SEM), B) Kaplan–Meier curves to 400mm^3^ tumor volume, and C) maximum loss of tumor volume for Rh5 cell line xenografts treated with erdafitinib 12 mg/kg (daily oral gavage for 21 days), FGF401 100 mg/kg (twice daily oral gavage for 21 days), lenvatinib 3 mg/kg (daily oral gavage for 21 days), or ponatinib 30 mg/kg (daily oral gavage for 21 days) with four mice per treatment group, six mice for vehicle control. **: P < 0.01 when compared to vehicle control by Mantel-Cox test. D) RNA TPM values for *FGFR4*, *FGF8* and *SPRY2* from *in vivo* tested FP-RMS and HGU sarcoma ZERO PDX models and the Rh5 cell line xenograft. E-H) Tumor growth (mean tumor volumes ± SEM) and I-L) Kaplan–Meier curves to 400 mm^3^ tumor volume plotted for zcc27 PDX (E,I), zcc31 (F,J) zcc81(G,K) and zcc275 (H,L) treated with FGF401 100 mg/kg (twice daily oral gavage for 26 days, zcc27 and zcc31 only) or lenvatinib 3 mg/kg (daily oral gavage for 26 days). *: P < 0.05, ***: P < 0.001, ****: P < 0.0001 when compared to vehicle control by Mantel-Cox test. M,N) Histopathology images of zcc27 (M) and zcc31 (N) FP-RMS PDX tumors harvested at end of treatment from vehicle control, FGF401 or lenvatinib-treated mice stained with H&E, cleaved caspase-3 or Ki67. Images captured at 600x.

The efficacy of FGF401 and lenvatinib was further examined in sarcoma PDX models of patients from ZERO. First, we tested sensitivities across predicted responders: zcc27 and zcc31 FP-RMS PDX models, harboring the *FGFR4+/FGF8+* expression signature (Figure 6D). In both FP-RMS zcc27 and zcc31 PDX models, lenvatinib and FGF401 therapies alone were effective in eliciting marked tumor growth delays (Figure 6E-F). Single agent FGF401 and lenvatinib also prolonged EFS compared to vehicle control (Figure 6I-J). Histological examination with H&E revealed that vehicle control, FGF401 and lenvatinib treated tumors from both models looked morphologically similar (Figure 6M-N). There were no apparent differences in the number of apoptotic cells, and only minimal necrosis (∼5%) was identified in lenvatinib-treated tumors from both models. The Ki67 proliferative index on the other hand was reduced upon FGFR-targeted treatments, most notably in FGF401-treated tumors (<5% Ki67-positive cells in zcc27; 10-20% in zcc31) compared to lenvatinib (40% Ki67-positive cells in zcc27; 20% in zcc31) and vehicle (40% both models). These findings are in keeping with a cytostatic rather than a cytotoxic drug effect.

As part of the ZERO preclinical testing program, where only clinically available drugs are tested in PDX models to guide individualized patient treatment recommendations, we next prospectively tested lenvatinib in a PDX model predicted to be sensitive on the basis of FGFR4 and FGF8 expression (FP-RMS zcc81; *FGFR4+/FGF8+*), and compared this to a sarcoma PDX model predicted to be resistant (HGU zcc275; *FGFR4-/FGF8-*)(Figure 6D). We note a sustained and impressive inhibition of tumor growth in the zcc81 FP-RMS PDX model, where none of the mice reached an event during the course of treatment (Figure 6G,K). In the predicted non-responding zcc275 model, all mice progressed eventually while on lenvatinib treatment (Figure 6H,L).

As lenvatinib is combined with etoposide and ifosfamide in OS patients, we tested the potential of this combination in FP-RMS [31]. *In vitro*, lenvatinib combined with etoposide and the active metabolite of ifosfamide (palifosfamide) resulted in additive anti-tumor effects, including more cell death in Rh5 FP-RMS cells treated with the combination compared to the highest tested concentrations of each drug alone (Supplemental Figure 11). We also tested the efficacy of this combination in sarcoma PDX models (Supplemental Figure 12). Here, we observed an unexpected sensitivity of most FP-RMS PDX models to clinically relevant doses of etoposide and ifosfamide. The HGU zcc275 PDX model, which was treated with the same regimen and dosing, did not respond to chemotherapy. The unexpected sensitivity of FP-RMS to this cytotoxic regimen precluded robust testing of the added value of chemotherapy combined with lenvatinib. We do however note that in an additional FP-RMS PDX model (zcc460), which we excluded from formal analysis due to variable growth of control tumors, substantial tumor shrinkage was observed in all mice treated with the lenvatinib-chemotherapy combination, including complete tumor regressions, which was superior to any other treatment arm (Supplemental Figure 12). Altogether, these data provide interesting hints of activity that warrant more research into the potential of such a combination in FP-RMS.

Based on our preclinical work, FGFR-inhibitors are now prospectively offered as an option (“recommended”) to FP-RMS patients in ZERO, based on the *FGFR4/FGF8* co-expression signature.

### Clinical response of FP-RMS patient to lenvatinib

The potential for TKi to impact on FP-RMS patients is illustrated in the following patient response in the ZERO cohort. A 15-year-old male, diagnosed with Stage 4 Group 4 FP-RMS and bony metastases (zcc153) was enrolled on ZERO and subsequently presented at the MTB. The *PAX3-FOXO1* gene fusion was confirmed and high expression of *FGFR4* was reported. Other reportable findings included high expression of *CDK4* and *CCND3*. Further analysis revealed this patients’ gene expression profile to be consistent with our proposed signature of sensitivity, with high *FGFR4* (520 TPM) and high *FGF8* (50 TPM), and as expected low *SPRY2* (73 TPM). We have subsequently confirmed activation of FGFR4 by pY MS analysis in the PDX model of this patient (Figure 3H).

The patient was treated with multiagent chemotherapy, and radiotherapy to the primary and metastatic masses. Surveillance imaging demonstrated multifocal perianal and retroperitoneal relapse of FP-RMS confirmed on FDG-PET scanning (Figure 7A). He was treated on a study of lenvatinib for patients with relapsed or refractory solid malignancies. The patient had a partial response to 4 cycles of lenvatinib (Figure 7B) but was then found to have a leptomeningeal relapse. After a 4 week pause of lenvatinib for radiotherapy, lenvatinib was recommenced. Follow up imaging demonstrated continued stability or improvement in all sites of disease (Figure 7C) although ultimately disease progression occurred at all sites.

**Figure 7.**
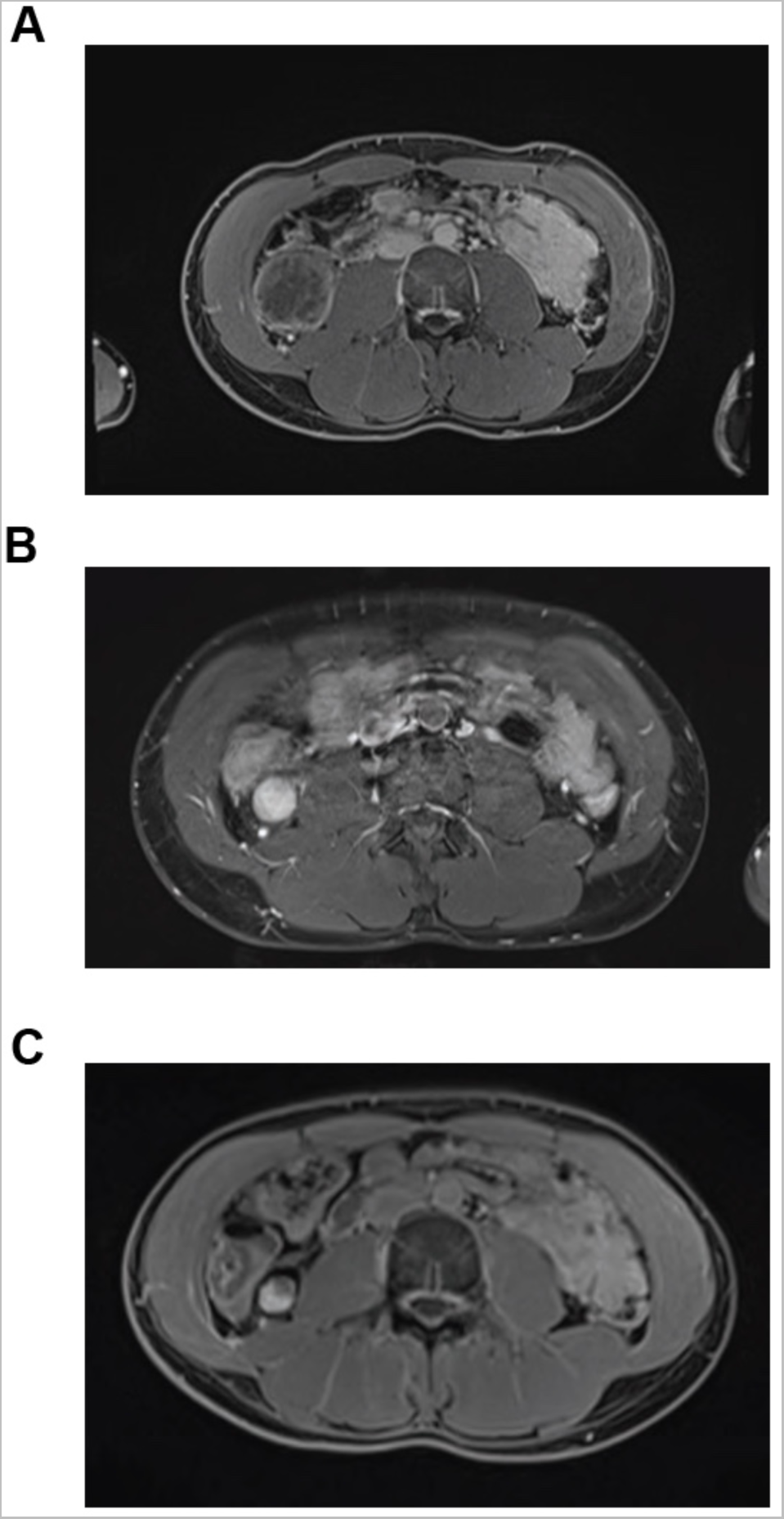
Lenvatinib treatment results in tumor regression and stabilization in a FP-RMS patient. Axial T1 fat saturation post contrast MRI images of FP-RMS patient zcc153. A) Baseline imaging of multifocal perianal and retroperitoneal disease relapse after chemotherapy treatment. Maximal diameter of right paracolic metastasis: 45 mm. B) Follow up imaging after four cycles of lenvatinib. Maximal diameter of right paracolic metastasis: 25 mm. C) Follow up imaging after seven cycles of lenvatinib. Maximal diameter of right paracolic metastasis: 18mm.

## Discussion

In this study, we integrated data from a comprehensive, multi-platform precision medicine program to paint a complete and unbiased picture of actionable TK aberrations in one of the largest and most diverse sarcoma cohorts of high-risk pediatric and AYA sarcoma patients investigated to date. Our approach coupled whole genomic and transcriptomic TK features in clinical sarcoma samples to phosphoproteomics and used functional *in vitro* and *in vivo* assays in sarcoma cell line models, isogenic model systems, and unique PDX models to validate newly identified therapeutically actionable aberrations in the FGFR-pathway. A key finding is that FGFRi may be beneficial to a subset of sarcoma patients who would currently not have such drugs included in their clinical treatment plans. This includes patients with novel, but rare activating fusion oncogenes such as *LSM1-FGFR1* as identified in an OS patient.

Our most significant finding is the actionability of the *FGFR4+/FGF8+* co-expression signature in FP-RMS, which may expand the therapeutic options for these patients. Activating mutations in *FGFR4* in RMS are rare (∼7.5-10%) [32], but the *FGFR4+/FGF8+* overexpression and associated FGFR4_pY activation signatures we report are recurrently observed across FP-RMS. This signature renders FP-RMS particularly vulnerable to FGFR inhibition in multiple *in vitro* and *in vivo* models. The apparent inhibitory effects observed with the clinically applicable multi-kinase inhibitor lenvatinib in our *in vivo* PDX models provides important preclinical data on the efficacy of lenvatinib alone, and its potential combination with chemotherapy. The clinical response, including partial tumor regression, to single agent lenvatinib in a relapsed FP-RMS patient harboring our molecular signature who exhausted all other treatments, provides a remarkable anecdote of the potential these drugs may have. Our findings are especially relevant in the field of advanced sarcoma, where tumors are notoriously difficult to treat and the overall prognosis is dismal. As a reference, pazopanib is the only approved targeted agent for metastatic soft-tissue sarcoma in adults (excluding GIST adipocytic sarcoma), and its approval was based on a significant gain in progression free survival of 3 months compared to placebo, without benefit to overall survival [33]. Although clinical trials in more FP-RMS patients are required, these data offer an exciting proposition for a tumor which, when metastatic, has a miserable prognosis.

The oncogenic role of TKs with elevated expression in the absence of a genomic aberration is controversial, and WT *FGFR4* expression is no exception. Several previous studies have investigated the role and targetability of FGFR4 in RMS, predominantly in the context of activating mutations but not in WT expression levels. In a murine model, overexpression of WT *FGFR4* did not contribute to tumorigenic transformation of primary mouse myoblasts, and knockdown of *FGFR4* actually enhanced growth in Rh4 cells [34]. In contrast, more recent work showed that overexpression of WT *FGFR4* in mouse myoblasts did result in the development of sarcomas in mice with high penetrance, although they lacked RMS-specific tumor markers [35]. shRNA knockdown further indicated that *FGFR4* was necessary for survival in FP-RMS Rh28 and Rh30 cells, as *FGFR4* suppression induced apoptotic cell death. This study also demonstrated that treatment of FP-RMS cells with PD173074 (an FGFR1 and VEGFR2 inhibitor with additional FGFR4-inhibitory effects) reduced Rh28 cell growth *in vitro,* and induced apoptosis in both Rh28 and Rh30 cells. PD173074 treatment also induced Rh28 tumor regression *in vivo,* albeit at toxic levels. The authors acknowledge that nonspecific toxicity may have influenced the study outcomes and concluded that pharmacologic blockade of FGFR4 will require alternative approaches [36]. Ponatinib at 30mg/kg daily further showed a lack of efficacy in RMS772 xenografts transduced with WT *FGFR4* [29].

These contradictions in the literature surrounding widespread applicability of FGFRi in FP-RMS have resulted in a reticence to move forward into clinical applications. This is reflected within ZERO; prior to this study, high *FGFR4* expression was not considered targetable in FP-RMS patients and thus did not lead to clinical recommendation of FGFRi. Whilst *FGFR4* can function as an oncogene, the important question is whether targeting *FGFR4* overexpression is a therapeutically valid approach. Our data argue that in some molecular contexts, WT *FGFR4* is a legitimate target, where we highlight its co-expression with *FGF8*.

The activation and actionability of WT *FGFR4* was recently also demonstrated in breast cancer models. FGFR4 protein overexpression and phosphorylation in a luminal B breast cancer PDX model conferred FGFR4-inhibitor sensitivity. The *FGFR4* gene was not amplified, suggesting a transcriptional or post-transcriptional mechanism of activation much like our findings [37]. In murine breast cancer models, a neutralizing anti-FGF8 antibody further showed potent effects *in vitro* and *in vivo.* This application has not been investigated in RMS [38].

An interesting question is whether *FGF8* is the only biomarker ligand indicating FGFR4 activation in FP-RMS. FGF19 is a well-known ligand of FGFR4, and its expression together with WT *FGFR4* and co-regulator *KLB*, was identified as a marker of FGF401-sensitive hepatocellular carcinoma [39–41], leading to the first-in-human Phase I/II study of FGF401 [40–42]. In our study, we found no evidence of *FGF19* or *KLB* upregulation in FP-RMS. There is evidence to indicate that FGF8 expression is induced by the canonical driver fusion of FP-RMS, *PAX3-FOXO1* [27]. Notably, both *FGF8* and *FGFR4* are dependency genes in FP-RMS [43]. We now comprehensively describe FGFRi sensitivity in FP-RMS that has not been previously established, but is consistent with the hypothesis that FGF8/FGFR4 signalling is a critical oncogenic signalling pathway in FP-RMS.

In addition to the FGFR-pathway, our comprehensive approach also shed light into the biology and potential actionability of a range of other TKs in sarcoma. We captured and mapped outlier levels of lesser-known components of the PDGFR-pathway, including *PDGFRL* and the ligands *PDGFC* and *PDGFD*, where we highlight overexpression in OS, a desmoid tumor and a FMX tumor. Further functional studies of the PDGFR-pathway and ligand dependency are warranted, focusing on combination strategies [44]. Our expression data also suggests that *ERBB3* and *FYN* might be implicated in the oncogenesis of selected sarcomas, and may hence represent actionable targets in individual patients, pending validation. We further showed that the cell surface receptor *PTK7* is recurrently expressed across different sarcoma subtypes, with outlier levels reported in a desmoid tumor patient. PTK7 CAR-T cells have been developed and may be applicable to treatment of some sarcomas after further research [45]. An important therapeutic question that flows from our work is which other TK RNA overexpression signatures in sarcoma are also actionable targets? These will not be recognized by genomic aberrations, but instead by functional preclinical studies, (phospho)proteomics and drug screens.

In conclusion, our studies provide a compelling rationale to implement screening of the complete genomic and transcriptomic landscape of TKs, to identify unrecognized drivers and expression signatures that can be therapeutically targeted in pediatric and AYA sarcomas. As a direct result of our work, the FGFRi lenvatinib, with or without chemotherapy, is now prospectively recommended as an option for FP-RMS patients in ZERO. This has already resulted in clinical benefit for a FP-RMS patient who exhausted all other treatment options. We therefore believe FGFR inhibitors represent a promising therapeutic avenue for FP-RMS patients, deserving further clinical investigation.

## Acknowledgements

We thank the Cancer Institute New South Wales (Fellowship funding for EDGF; 2020/ECF1261) and the Can Too Foundation (Emerging Research Leader Program Grant funding for EDGF) for supporting this study. We sincerely thank patients and parents for participating in this study. We thank the many clinicians and health professionals for their time acquiring consenting patients and for collection and coordination of samples and associated clinical data at Sydney Children’s Hospital, Randwick; The Children’s Hospital at Westmead; the John Hunter Hospital; the Queensland Children’s Hospital; the Royal Children’s Hospital Melbourne; the Monash Children’s Hospital; the Adelaide Women’s & Children’s Hospital; and the Perth Children’s Hospital.

We also thank the Sydney Children’s Tumor Bank Network (Children’s Cancer Institute (CCI) Tumor Bank and/or Children’s Hospital at Westmead Tumor Bank) for providing samples and related clinical information for this study. We thank Tour de Cure for supporting CCI Tumor Bank personnel and M. Weber (Prince of Wales Hospital) for assistance with anatomical pathology expert input.

We thank the staff of the Personalized Medicine Theme of the Children’s Cancer Institute for their dedicated work on ZERO Childhood Cancer.

We thank the staff at the Murdoch Children’s Research Institute/Victorian Comprehensive Genetics Service for processing of RNAseq.

We thank the team from the National Computational Infrastructure, which is supported by the Australian Government. We thank the Australian Federal Government Department of Health, the New South Wales State Government and the Australian Cancer Research Foundation for funding to establish infrastructure to support the ZERO Childhood Cancer personalized medicine program. We thank the Kids Cancer Alliance, Cancer Therapeutics Cooperative Research Centre, for supporting the development of a personalized medicine program; Tour de Cure for supporting tumor biobank personnel; Samuel Nissen Charitable Foundation “Translating Molecular Discoveries in Childhood Cancer into Improved Outcomes” for supporting P.G.E.; and the Lions Kids Cancer Genome Project, a joint initiative of Lions International Foundation, the Australian Lions Children’s Cancer Research Foundation (ALCCRF), the Garvan Institute of Medical Research, the Children’s Cancer Institute and the Kids Cancer Centre, Sydney Children’s Hospital. Lions International and ALCCRF provided funding to perform WGS and for key personnel. We thank the Kinghorn Foundation for personnel support. We thank the Kids Cancer Project for supporting molecular profiling and molecular and clinical trial personnel; and the University of New South Wales, W. Peters and the Australian Genomics Health Alliance for providing personnel funding support.

The authors would like to acknowledge Luminesce Alliance - Innovation for Children’s Health for its contribution and support. Luminesce Alliance - Innovation for Children’s Health, is a not-for-profit cooperative joint venture between the Sydney Children’s Hospitals Network, the Children’s Medical Research Institute, and the Children’s Cancer Institute. It has been established with the support of the NSW Government to coordinate and integrate pediatric research. Luminesce Alliance is also affiliated with the University of Sydney and the University of New South Wales Sydney. The Medical Research Future Fund, the Australian Brain Cancer Mission, Minderoo Foundation’s Collaborate Against Cancer Initiative and funds raised through the ZERO Childhood Cancer Capacity Campaign, a joint initiative of the Children’s Cancer Institute and the Sydney Children’s Hospital Foundation, supported the national clinical trial and associated clinical and research personnel.

We thank the National Health and Medical Research Council of Australia (fellowship APP1157871 to R.B.L), the Cancer Institute of New South Wales and New South Wales Health (fellowship funding for M.J.C.). We thank the 2018 Priority-Driven Collaborative Cancer

Research Scheme, co-funded by Cancer Australia and My Room, for personnel and computational support (grant no. 1165556 awarded to M.J.C.).

We also sincerely thank Peter Houghton (Greehey Children’s Cancer Research Institute, USA; Ewing- and rhabdomyosarcoma cell lines), Kazuyuki Itoh and Norifumi Naka (Osaka Medical Center for Cancer and Cardiovascular Diseases, Japan; Yamato-SS and Aska-SS), Akira Kawai (National Cancer Center Hospital, Japan; SYO-1), Cinzia Lanzi (Fondazione IRCCS Istituto Nazionale dei Tumori, Italy; CME-1), Mikio Masuzawa (Kitasato University School of Allied Sciences, Japan; MO-LAS-B and ISO-HAS-B) and Jonathan Fletcher (Brigham and Women’s Hospital, Harvard Medical School, USA; GIST882) for kindly providing sarcoma cell lines.

ZERO Childhood Cancer is a joint initiative led by the Children’s Cancer Institute and Sydney Children’s Hospital, Randwick.

