## Supplemental Data_Legends and Methods for "Comprehensive multi-platform tyrosine kinase profiling reveals novel actionable FGFR aberrations across pediatric and AYA sarcomas"

### Supplemental Table Legends

**Supplemental Table 1. Cell culture conditions**
Cell lines were cultured in optimized conditions. AS: angiosarcoma, EWS: Ewing sarcoma, GIST: gastrointestinal stromal tumor, OS: osteosarcoma, FP-RMS: fusion-positive rhabdomyosarcoma, FN-RMS: fusion-negative rhabdomyosarcoma, SynS: synovial sarcoma.

**Supplemental Table 2. Pediatric and AYA sarcoma patient cohort characteristics**
Cohort of 107 pediatric and AYA sarcoma patients from the ZERO Childhood Cancer program included in this study ("sarcoma cohort"); reportable data shown (see Methods). Characteristics, including patient identifier number (zcc***), diagnosis, disease stage, age at diagnosis, age at sampling, vital status, gender and reportable tyrosine kinase (TK) molecular aberrations (SNV: single nucleotide variation, CNV: copy number variation, SV: structural variation, or somatic RNA (Z-score, calculated from ZERO cohort at time of analysis) in the tumor and germline, along with tyrosine kinase inhibitor (TKi) recommendations made to the ZERO multidisciplinary tumor board (MTB) based on these molecular TK features, are recorded. Recommendations are assigned a tier according to the following criteria: Tier 1, clinical evidence in the same cancer; Tier 2, clinical evidence in a different cancer; Tier 3, preclinical evidence in the same cancer; Tier 4, preclinical evidence in a different cancer type; Tier 5, consensus opinion [1].

**Supplemental Table 3. Tyrosine kinase RNA expression in the pediatric and AYA sarcoma cohort**
Cohort of 86 pediatric and AYA sarcoma patients, and two patients with other soft-tissue malignancies (one desmoid and one osteoblastoma) from the ZERO Childhood Cancer program; tyrosine kinase (TK) RNA expression data shown. Values depicted as TPM counts.
EWS: Ewing sarcoma, FP-RMS: fusion-positive rhabdomyosarcoma, FN-RMS: fusion-negative rhabdomyosarcoma, OS: osteosarcoma, MRT: malignant rhabdoid tumor, MPNST: malignant peripheral nerve sheath tumor, ASPS: alveolar soft part sarcoma, AS: angiosarcoma, IFS: infantile fibrosarcoma, DSRCT: desmoplastic small round blue cell tumor, GIST: gastrointestinal stromal tumor, SynS: synovial sarcoma, GNET: gastrointestinal neuroectodermal tumor, HGU: high-grade undifferentiated sarcoma, AF: ameloblastic fibrosarcoma, CIC: CIC-rearranged sarcoma, EPI: epithelioid sarcoma, FMX: low-grade fibromyxoid sarcoma, Round cell: *EWSR1-PATZ1* round cell sarcoma, desmoid: desmoid tumor.

**Supplemental Table 4. FGFR pathway RNA expression in the pediatric and AYA sarcoma cohort**
Cohort of 86 pediatric and AYA sarcoma patients from the ZERO Childhood Cancer program; FGFR-pathway RNA expression data shown. Values depicted as TPM counts. Abbreviations as per Supplemental Table 3.

**Supplemental Table 5. FGFR pathway RNA expression in the sarcoma cell line panel**
Sarcoma cell line panel; FGFR-pathway RNA expression data shown. Values depicted as TPM counts.
EWS: Ewing sarcoma, FP-RMS: fusion-positive rhabdomyosarcoma, FN-RMS: fusion-negative rhabdomyosarcoma, OS: osteosarcoma, SynS: synovial sarcoma, GIST: gastrointestinal stromal tumor, AS: angiosarcoma.

**Supplemental Figure Legends**

**Supplemental Figure 1.**
A) PCR amplification of the *LSM1-FGFR1* fusion and breakpoint sequence identified in an osteosarcoma patient (zcc386). B) Viability analysis of *LSM1-FGFR1*- and *FGFR1OP2-FGFR1 (OP2-FGFR1)*-expressing Ba/F3 cells following 96-hour IL-3 withdrawal. Viability was determined by propidium iodide exclusion, measured by flow cytometry. Data presented as mean ± SEM ($n=3$). C) Viability analysis of *LSM1-FGFR1*- and *OP2-FGFR1*-expressing Ba/F3 cells treated with a dose titration of regorafenib, AZD4547 or imatinib. Data presented as mean ± SEM ($n=6$).

**Supplemental Figure 2.**
A) Heatmap visualizing RNA expression (TPM) of TKs across the sarcoma cohort ordered based on sarcoma histology. RNA expression (Z-score) for B) *JAK1*, C) *PDGFRB* or D) *PDGFRL* in the ZERO cohort. Colors indicate specific samples as shown in legend.
EWS: Ewing sarcoma, FP-RMS: fusion-positive rhabdomyosarcoma, FN-RMS: fusion-negative rhabdomyosarcoma, OS: osteosarcoma, MRT: malignant rhabdoid tumor, MPNST: malignant peripheral nerve sheath tumor, ASPS: alveolar soft part sarcoma, AS: angiosarcoma, IFS: infantile fibrosarcoma, DSRCT: desmoplastic small round blue cell tumor, GIST: gastrointestinal stromal tumor, SynS: synovial sarcoma, GNET: gastrointestinal neuroectodermal tumor, HGU: high-grade undifferentiated sarcoma, AF: ameloblastic fibrosarcoma, CIC: CIC-rearranged sarcoma, EPI: epithelioid sarcoma, FMX: low-grade fibromyxoid sarcoma, Round cell: *EWSR1-PATZ1* round cell sarcoma, desmoid: desmoid tumor.

**Supplemental Figure 3.**
A) *PDGFA-D* ligand RNA expression (TPM) across sarcoma samples, rank ordered on total PDGF-expression counts. RNA expression (Z-score) for B) *PTK7*, C) *FGFR4,* D) *FGFBP1-3* and E) *FGF8* in the ZERO cohort. Colors indicate specific samples as shown in legends. EWS: Ewing sarcoma, FP-RMS: fusion-positive rhabdomyosarcoma, FN-RMS: fusion-negative rhabdomyosarcoma, OS: osteosarcoma, MRT: malignant rhabdoid tumor, MPNST: malignant peripheral nerve sheath tumor, ASPS: alveolar soft part sarcoma, AS: angiosarcoma, IFS: infantile fibrosarcoma, DSRCT: desmoplastic small round blue cell tumor, GIST: gastrointestinal stromal tumor, SynS: synovial sarcoma, GNET: gastrointestinal neuroectodermal tumor, HGU: high-grade undifferentiated sarcoma, AF: ameloblastic fibrosarcoma, CIC: CIC-rearranged sarcoma, EPI: epithelioid sarcoma, FMX: low-grade fibromyxoid sarcoma, Round cell: *EWSR1-PATZ1* round cell sarcoma, desmoid: desmoid tumor.

**Supplemental Figure 4.**
A) Heatmap visualizing RNA TPM counts of the FGFR-pathway across the sarcoma cell line panel, ordered based on histology and FGFR genes. B) Correlation between FGFR4 tyrosine phosphorylation (sum of all pY sites; Figure 3E) and FGF8 protein expression (Figure 3C) from RMS cell lines.

**Supplemental Figure 5.**
A) Area under the curve (AUC) analysis for resazurin growth-inhibition assay after 76-hour inhibitor treatment with erdafitinib or ponatinib, and dose-response curve and AUC analysis for AZD4547. An *OP2-FGFR1* overexpression model was included as a positive control for non-FGFR4 specific inhibitors. ES8, EW8 and MG63 were included as negative, non-RMS, control sarcoma cell lines. Data presented as mean viable cells remaining or AUC ± SEM (*n=3*). B) Representative images of colony assays (*n=3* from Figure 4D-E) following two-weeks inhibitor treatment with lenvatinib or erdafitinib. C) Flow cytometry-based Annexin-V/7-AAD analysis. FP-RMS cell lines cells were harvested following 96-hour treatment with AZD4547 or ponatinib and stained with Annexin-V and 7-AAD. Gating was performed against positive controls for live (Annexin-V and 7-AAD negative), apoptotic (Annexin-V positive) or necrotic (7-AAD positive and Annexin-V negative) cells (gating strategy example: Supplemental Figure 5D) and applied to cells which were untreated or treated with FGFR4 inhibitors. Data presented as mean percentage of live, apoptotic or necrotic cells present in the total population ± SEM (*n=3*). D) Gating was performed against positive controls for live, apoptotic (Annexin-V positive or necrotic (7-AAD positive and Annexin-V negative) cells and applied to cells which were untreated or treated. Example plots demonstrating application of gating strategy to 0.1 μM and 1 μM AZD4547 treated Rh5 cells.

**Supplemental Figure 6.**
Images captured of Rh3 FP-RMS cells at 76h after treatment with serial dilutions of AZD4547, erdafitinib, FGF401, lenvatinib or ponatinib, at 10x magnification, unfiltered.

**Supplemental Figure 7.**
Images captured of Rh5 FP-RMS cells at 76h after treatment with serial dilutions of AZD4547, erdafitinib, FGF401, lenvatinib or ponatinib, at 10x magnification, unfiltered.

**Supplemental Figure 8.**
Images captured of Rh41 FP-RMS cells at 76h after treatment with serial dilutions of AZD4547, erdafitinib, FGF401, lenvatinib or ponatinib, at 10x magnification, unfiltered.

**Supplemental Figure 9.**
Images captured of Rh30 FP-RMS cells at 76h after treatment with serial dilutions of AZD4547, erdafitinib, FGF401, lenvatinib or ponatinib, at 10x magnification, unfiltered.

**Supplemental Figure 10.**
A) Over a two-week conditioning period with or without FGF8 at 20 ng/mL, total number of viable cells determined by trypan blue exclusion. Data presented as population doublings (PDL, see Supplemental Methods). B) Western blot analysis of Rh5 cells transduced with Cas9 and a doxycycline inducible guide RNA targeting FGF8 or controls (non-targeting gRNA or empty vector (Empty)) to confirm FGF8 knockout model. Doxycycline (+dox), or water control (-dox), was added to cells for 72 hours to induce gRNA expression and FGF8 was analysed after a further two days in culture. C-D) Effect of FGF8 KO out on cell proliferation determined by C) PDL growth assay over a five-week period or D) colony-forming assay over a two-week period for Rh5 cells with or without FGF8 KO. E) Clone 7 Rh30 SPRY2 knockout was used for resazurin growth-inhibition assays following 76-hour inhibitor treatment with lenvatinib, erdafitinib or FGF401. Data presented as mean viable cells remaining ± SEM (*n=3*). F) Two SPRY2 overexpression Rh5 models were used for resazurin growth-inhibition assays following 76-hour inhibitor treatment with lenvatinib, erdafitinib or FGF401. Data presented as mean viable cells remaining ± SEM (*n=3*)

**Supplemental Figure 11.**

Cell viability in the Rh5 FP-RMS cell line was measured 72 hours following treatment with increasing concentrations of A) lenvatinib in combination with increasing concentrations of a chemotherapy backbone (etoposide and palifosfamide), B) etoposide and palifosfamide with increasing concentrations of lenvatinib. Data presented as mean viable cells remaining ± SEM (*n=3*). C) Heatmap representation of percentage inhibition of Rh5 cells treated with varying concentrations of lenvatinib and etoposide-palifosfamide. D) Bliss synergy score was calculated, where a value >10 indicates synergy, a value between -10 to 10 indicates an additive effect and a value <-10 indicates an antagonistic effect.

**Supplemental Figure 12.**

Tumor growth (mean tumor volumes ± SEM), Kaplan–Meier curves (event defined as tumor exceeding 400 $mm^3$ volume) and maximum loss in tumor volume were plotted for FP-RMS zcc27 (A-C), FP-RMS zcc31 (D-F), FP-RMS zcc81(G-I), FP-RMS zcc490 (J-L) and HGU zcc275 (M-O) PDX models treated with FGF401 100 mg/kg (twice daily oral gavage for 26 days, zcc27 and zcc31 only), lenvatinib 3mg/kg (daily oral gavage for 26 days, all models), ifosfamide 90 mg/kg and etoposide 6 mg/kg (once daily intraperitoneal injection on days 1-5 and 22-26, all models) or lenvatinib 3 mg/kg (daily oral gavage for 26 days, all models) and ifosfamide 90 mg/kg and etoposide 6 mg/kg (once daily intraperitoneal injection on days 1-5 and 22-26, all models). *: $P < 0.05$, ***: $P < 0.001$, ****: $P < 0.0001$ when compared to vehicle control by Mantel-Cox test.

**Supplemental Methods**

**Patients and samples**

The pilot study (TARGET) was approved by the Sydney Children's Hospitals Network Human Research Ethics Committee (LNR/14/SCH/497) and opened at Sydney Children's Hospital, Randwick and The Children's Hospital at Westmead, from June 2015 to October 2017. The PRISM clinical trial (NCT03336931), conducted as part of ZERO, was approved by the Hunter New England Human Research Ethics Committee of the Hunter New England Local Health District (reference no. 17/02/015/4.06) and the New South Wales Human Research Ethics Committee (reference no. HREC/17/HNE/29). ZERO was opened at Sydney Children's Hospital Randwick, Sydney; The Children's Hospital at Westmead, Sydney; John Hunter Hospital, Newcastle; Queensland Children's Hospital, Brisbane; Royal Children's Hospital, Melbourne; Monash Children's Hospital, Melbourne; Women's & Children's Hospital, Adelaide; and Perth Children's Hospital, Perth, from September 2017 onwards. Parents/legal guardians for participants under the age of 18 years or by participants who were over the age of 18 years provided informed consent for study inclusion. Patients aged 21 years or younger with suspected or confirmed diagnosis of a very rare or high-risk malignancy (at diagnosis, relapse or refractory disease), defined as expected probability of survival of less than 30%, could be consented and were eligible for registration on the study. Registration of patients older than 21 years was possible with approval from the study chair when the patient was diagnosed with suspected or confirmed high-risk pediatric-type cancers. Following registration, patient tumor samples were delivered to the Children's Cancer Institute in Sydney. Study enrollment was subject to clinical confirmation of high-risk cancer diagnosis and receipt of both a tumor and a germline sample at the Children's Cancer Institute. Tumor tissue (solid tissue or tumor cells isolated from bone marrow aspirate or peripheral blood) was fresh, fresh frozen or cryopreserved upon receipt. When patients were recruited at relapse or after the onset of refractory disease, retrospective access to diagnostic samples was not sought. In the case of patients who had undergone a bone marrow transplantation during their treatment, both the patient germline and the donor germline were sequenced specifically to distinguish tumor-derived somatic variants from donor variants.

**Overexpression model generation (*LSM1-FGFR1*, *SPRY2*)**

For generation of the LSM1-FGFR1 construct, patient RNA was reverse transcribed into cDNA using the SuperScript™ III First-Strand Synthesis System (Thermo Fisher Scientific, Cat#18080051) under standard conditions. *LSM1-FGFR1* full-length cDNA was PCR amplified from patient cDNA using AmpliTaq Gold™ DNA Polymerase with Buffer II and $MgCl_2$ (Thermo Fisher Scientific, Cat# N8080241) using the following primers: *LSM1* FL For: 5'-ATGAACTATATGCCTGGC and *FGFR1* FL Rev: 5' ACAGGGACGGACAGGTGG according to Manufacturer's instructions. Full-length *LSM1-FGFR1* cDNA was then subcloned into the MSCV-IRES-GFP retroviral vector using standard procedures – restriction digest, ligation of the digested product into MSCV-IRES-GFP using T4 DNA ligase (New England Biolabs, Cat# M0202L), and transformation of NEB Stable cells (New England Biolabs, Cat# C3040H). For generation of SPRY2 constructs, Rh30 RNA was reverse transcribed into cDNA and SPRY2 cDNA was PCR amplified as described previously, using the following primers: BamHI SPRY2 forward: 5'-TAG<u>GGATCC</u>ATGGAGGCCAGAGCTCAG, and EcoRI SPRY2 reverse: 5'-TCG<u>GAATTC</u>CTATGTTGGTTTTTCAAAG under standard conditions. SPRY2 cDNA was cloned into the doxycycline-inducible pFTRE tight mTAad GFP (pFTRE) lentiviral vector using standard procedures. Two SPRY2 variants were amplified from Rh30 cDNA, wild-type SPRY2 (referred to as "SPRY2#1") and P106S SPRY2 (referred to as "SPRY2#2", representing a SPRY2 variant observed in the normal population and predicted to have no functional effect), and subsequently cloned into pFTRE for functional studies. Retroviral and lentiviral transfection and transduction of target cell lines with respective plasmids, and empty vector controls, was performed as previously described [2]. Flow cytometry analysis of transduced cell populations was used to confirm >90% GFP positive cells or cells were sorted by Fluorescence-Activated Cell Sorting (UNSW Flow Cytometry Facility, UNSW Sydney) before proceeding to functional assays.

**CRISPR/Cas9 gRNA cloning and knockout model generation**

For FGF8 knockouts, a commercially validated sgRNA sequence (Horizon Bioscience) was selected and cloned into the doxycycline inducible gRNA vector with GFP reporter FgH1tUTG [3]. For SPRY2 knockouts, guide RNAs (gRNAs) targeting human *SPRY2* were designed using the CHOPCHOP web-based tool for CRISPR/Cas9 guide design [4]. The top two ranked guides (SPRY2 sgRNA#1 and SPRY2 sgRNA#2) were selected and cloned into FgH1tUTG. For FGF8 studies, Rh5 and Rh41 cells were transduced with FUCas9Cherry and FgH1tUTG containing FGF8 gRNA or non-targeting gRNA. Rh30 cells were transduced with FUCas9Cherry and FgH1tUTG containing SPRY2 sgRNA#1, SPRY2 sgRNA#2 or no sgRNA (control) lentiviruses, as previously described. A double positive population of >90% was confirmed by flow cytometry analysis of GFP and mCherry expression in transduced cells. Guide expression was induced by the addition of 1µg/mL doxycycline (Sigma Aldrich, Cat# D9891-1G) for 72 hours. Following guide induction, double stranded breaks have been induced and doxycycline is washed out of the media. Bulk CRISPR edited populations were analyzed by western blot to assess guide efficiency at least 4 days after doxycycline induction. For SPRY2, sgRNA#1 was determined to be the most efficient guide at inducing knockout in bulk cell populations, and these cell populations were subjected to single cell sorting (UNSW Flow Cytometry Facility) to generate single cell clonal populations. CRISPR-edited single cell SPRY2 knockout clone #7 was subjected to functional analysis.

**Western Blot**

Protein extraction was performed by the addition of RIPA lysis buffer containing cOmplete$^{TM}$ protease inhibitor cocktail (Roche, Cat#11697498001) and PhosSTOP (Roche, Cat#4906845001) phosphatase inhibitor to cell pellets followed by disruption by vortexing. PDX tumor lysates were generated by homogenization of a snap frozen tumor piece in RIPA buffer with protease inhibitor cocktail and PhosStop. Protein quantification was performed by Pierce$^{TM}$ BCA assay (Thermo Fisher Scientific, Cat#23225). Membranes were incubated with the following primary antibodies diluted in 1% bovine serum albumin (BSA) (Sigma-Aldrich, Cat# A7906) or 5% skim milk in Tris base sodium chloride (Merck, Cat# T6664) with 0.1% Tween-20 (Sigma-Aldrich, Cat#11332465001) (TBST) and incubated overnight at 4 °C: anti-p-FGFR1 antibody (milk, Abcam, Cat#Ab173305) at 1:1000, anti-FGFR1 antibody (milk, Cell Signaling, Cat#9740) at 1:5000, anti-p-AKT antibody (BSA, Cell Signaling, Cat#4060) at 1:500, anti-AKT antibody (milk, Cell Signaling, Cat#4691) at 1:500, anti-p-ERK1/2 antibody (BSA, Cell Signaling, Cat#9106S) at 1:500, anti-ERK1/2 antibody (BSA, Cell Signaling, Cat#4695) at 1:500, anti-FGF8 (milk, AbCam, Cat#81384) at 1:250, anti-FGFR4 (BSA, Santa-Cruz, Cat# 136988) at 1:100, anti-SPRY2 (milk, Cell Signaling, Cat#14954) at 1:500, or anti-actin antibody (milk, Sigma-Aldrich, Cat#A2228) at 1:1000. HRP-conjugated secondary donkey anti-rabbit (GE Healthcare, Cat# GEHENA934) or sheep anti-mouse (GE Healthcare, Cat# NA931) were diluted 1:1000 in 1% BSA or 5% milk in TBST and incubated at room temperature for 2 h.

***In vitro* growth-inhibition assays**
Cells were seeded into 96-well plates in a final volume of 100μL (sarcoma cell lines) or 200uL (Ba/F3 cells) of normal culture medium. Rh3 cells were seeded at $8 \times 10^3$ cells/well, Rh5, Rh36, Rh41, M25FV24C cells at $6 \times 10^3$ cells/well, RD and EW8 cells at $4 \times 10^3$ cells/well, and Rh30, ES8 and MG63 cells at $3 \times 10^3$ cells/well. Ba/F3 cells were seeded at $1 \times 10^5$ cells/mL or $3 \times 10^5$ cells/ml in 96-well round-bottom plates. For the FGF8 experiments, cell lines were pre-conditioned for two-weeks in 20 ng/mL FGF8 prior to seeding and seeding and treatment were also performed with FGF8 supplementation. PDL calculated by Log10(total number of cells counted ÷ number of cells seeded) ÷ log10(2). Serial dilutions of inhibitors (Selleck Chemicals) erdafitinib (Cat#S8401), lenvatinib (Cat#S5240) or FGF401 (Cat#S8548), AZD4547 (Cat#S2801), regorafenib (Cat#S1178), imatinib (Cat#S2475) or ponatinib (Cat#S1490), or 10 μM Thonzonium Bromide (100% cell death control, Focus Bioscience, Cat#HY-B1246) were added to growth media, in the presence or absence of doxycycline (1 μg/mL, Sigma-Aldrich, Cat#D9891) or FGF-8a (10ng/mL, Peprotec Cat#100-25A) and FGF-8b (10ng/mL, Peprotec Cat#100-25) and incubated under normal culture conditions of 5% CO2 for 48 or 96 h prior to assessment by resazurin assay. For assays examining response in the presence of FGF8, cells were pre-conditioned with FGF8 for 14 days (or more) prior to seeding. Data is normalized to vehicle control (DMSO). Responses were analyzed by four-parameter logistic model by non-linear regression and area under the curve in GraphPad Prism 9.2.0, with statistical analysis by one-way ANOVA with Tukey's multiple comparisons test (GraphPad Software).

**Mass Spectrometry**
Cell pellets or PDX snap frozen pieces were lysed in 4% sodium deoxycholate at 95°C for 5 min. Following sonication, protein concentration was determined by BCA assay. 1.5 mg of protein was used as starting material. For reduction and alkylation 10 mM TCEP and 40 mM 2-chloroacetamide (pH 7-8) was added to the lysates and boiled at 95°C for 5 min. Lysates were then digested with LysC and Trypsin at an enzyme-to-substrate ratio of 1:100 and incubated overnight at 37°C with shaking at 1500 rpm.

The rest of the protein was used for phospho-tyrosine enrichment by affinity purification, as previously described [5]. 1 mg of protein digest was acidified to 1%TFA and the resulting cleared supernatant was desalted using Sep-Pak tC18 column. The peptides were eluted with 6 mL 0.1% TFA/40% ACN, dried overnight using a lyophiliser and reconstituted in 1.8 mL IAP wash buffer (1% n-octyl-b-D-glucopyranoside, 50 mM Tris-HCl, 150 mM NaCl, pH 7.4). 50 μg each of P-Tyr-1000 (Cell Signaling Technology, Cat#8954), P-Tyr-100 (Cell Signaling Technology, Cat#9411), and P-Tyr-20 (BD Biosciences, Cat#610000) antibodies were coupled to 60 μL of protein G sepharose beads slurry (Rec-Protein G, Zymed) and incubated overnight with peptide samples at 4°C with gentle shaking. The conjugated antibody beads were washed three times with IAP buffer and further washed three times with water before phosphopeptide elution with 100 μL of 0.15% TFA. Samples were then desalted on a C18 column and evaporated to dryness in a vacuum concentrator (CentriVap™ Labconco).

Samples were analysed on an UltiMate 3000 RSLC nano LC system (Thermo Fisher Scientific) coupled to a Q Exactive HF Mass Spectrometer (Thermo Fisher Scientific). The samples were injected onto a 100 μm, 2 cm nanoviper Pepmap100 trap column, eluted and separation performed on a RSLC nano column 75 μm x 50 cm, Pepmap100 C18 analytical column (Thermo Fisher Scientific). Peptides were eluted using an LC gradient of 2.5-42.5% ACN/0.1%FA at 250 nL/min over 150 min. The mass spectrometer was operated in the data-dependent acquisition mode to automatically switch between full MS scans and subsequent

MS/MS acquisitions. Survey full scan MS spectra (m/z 300–1750) were acquired in the Orbitrap with 70,000 resolution (at m/z 200) after accumulation of ions to a $1x10^6$ target value with a maximum injection time of 30 ms. Dynamic exclusion was set to 30 s. Up to ten most intense charged ions ($z \geq +2$) were sequentially isolated and fragmented in the collision cell by higher-energy collisional dissociation with a fixed injection time of 120 ms, 17,500 resolution and automatic gain control target of $1x10^5$. The raw MS files were processed using MaxQuant software (v6.12.0) with the following parameters: FDR <0.01, precursor mass tolerance set to 20 ppm, fragment mass tolerance set to 0.5 Da, minimum peptide length of six amino acids, enzyme specificity set to Trypsin/P & LysC, human uniport database (v2020) and a maximum number of missed cleavages of 2. Fixed modifications were limited to Carbamidomethyl (C) and variable modifications were set to Oxidation (M), Acetyl (protein N-term), and Phospho (STY). The 'match between runs' option in MaxQuant was selected using the default parameters.

**Colony Assays**
Seeding densities for colony assays were as follows; Rh3 $1x10^4$ cells/well, Rh5 $0.2x10^4$ cells/well, Rh41 $0.4x10^4$ cells/well, ES8 $0.5x10^4$ cells/well in 3 mL culture medium.

***In vivo* drug efficacy studies in sarcoma xenografts**
Doses selected for animal studies are, to the best of our knowledge, reflecting clinically achievable doses. Erdafitinib was administered at 12 mg/kg (daily oral gavage for 21 days (Rh5 cell model)), FGF401 at 100 mg/kg (twice daily oral gavage for 21 (Rh5 cell model) or 26 (PDX models) days), lenvatinib at 3 mg/kg (daily oral gavage for 21 (Rh5 cell model) or 26 (PDX models) days), ponatinib at 30 mg/kg (daily oral gavage for 21 days (Rh5 model)). The combination of ifosfamide and etoposide for all PDX models was as follows: ifosfamide 90 mg/kg and etoposide 6 mg/kg (once daily intraperitoneal injection on days 1-5 and 22-26). We paid special attention to the dosing regimen associated with the active clinical trial for pediatric and adolescent osteosarcoma patients (NCT02432274), where lenvatinib is combined with etoposide and ifosfamide. In this trial, lenvatinib was administered orally, once daily on days 1 to 21 of each 21-day cycle at 8.8 $mg/m^2$. A 3 mg/kg dose in mice is calculated to be equivalent to 9 $mg/m^2$ per dose. Ifosfamide was administered at 3000 $mg/m^2$/day (starting dose) on days one to three of each 21-day cycle for a total of five cycles. Ifosfamide dose can be de-escalated to 2400 $mg/m^2$/day and 1800 $mg/m^2$/day. The maximum total ifosfamide dose was 9000 $mg/m^2$ per three-day cycle. The dose administered to mice in this study is calculated to be equivalent to 270 $mg/m^2$ per dose, a total of 1350 $mg/m^2$ in the first cycle of five days, thus below the achievable patient dose. Finally, etoposide 100 $mg/m^2$/day (starting dose) was administered on days one to three of each 21-day cycle for a total of five cycles. Etoposide dose can be de-escalated to 80 $mg/m^2$/day and 60 $mg/m^2$/day (total = 300 in the first cycle of three days). The dose administered to mice in this study is calculated to be equivalent to 18 $mg/m^2$ per dose (total is 90 $mg/m^2$ in the first cycle of five days), thus below the achievable patient dose [6].
