## Supplemental Figures for "Comprehensive multi-platform tyrosine kinase profiling reveals novel actionable FGFR aberrations across pediatric and AYA sarcomas"

#### Supplemental Figure 1

A

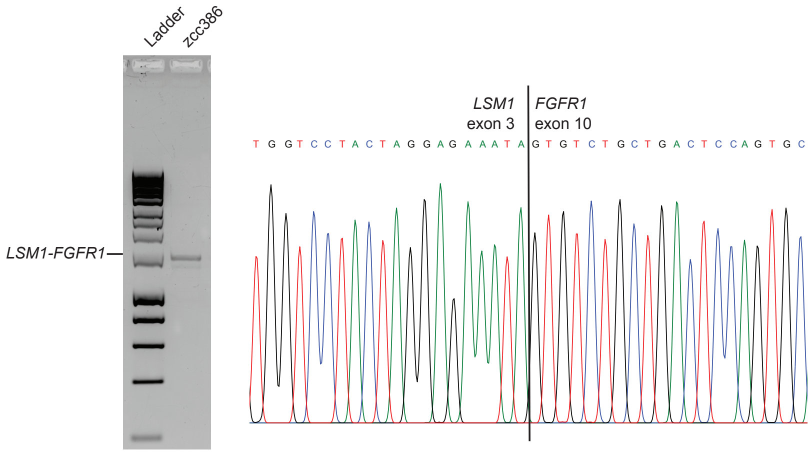


B

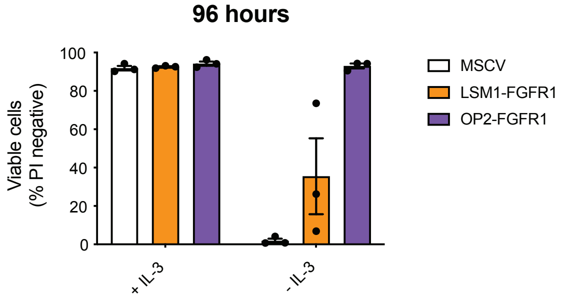


C

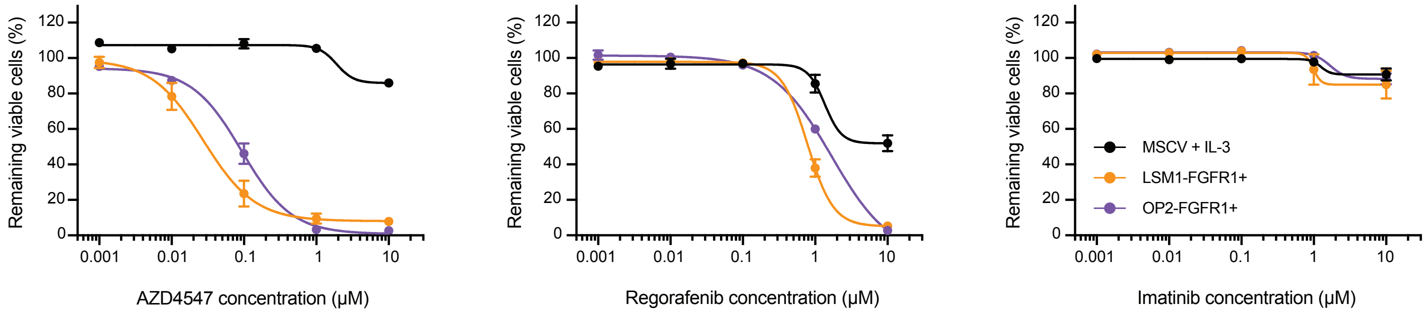

#### Supplemental Figure 2

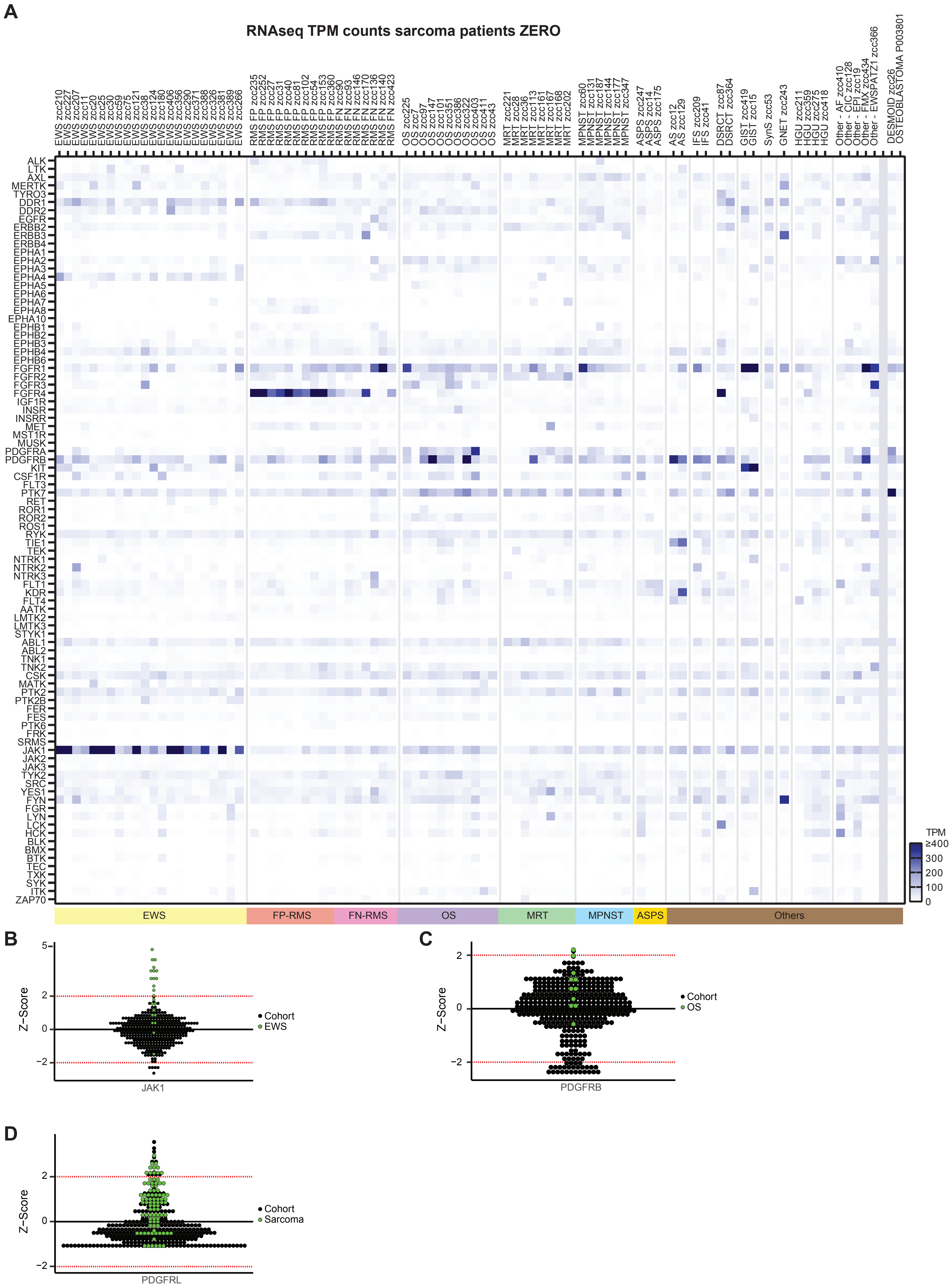

#### Supplemental Figure 3

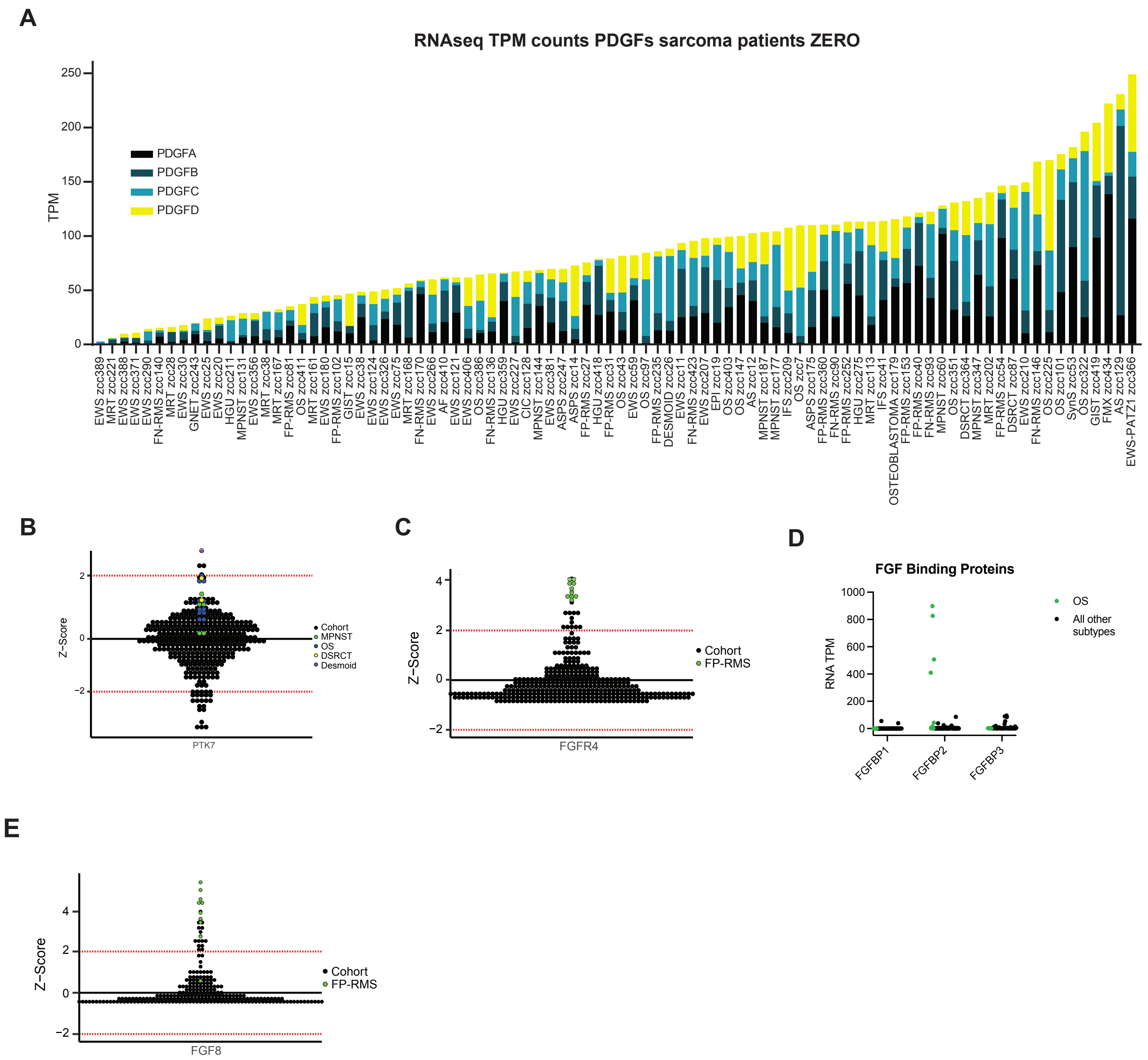

#### Supplemental Figure 4

A

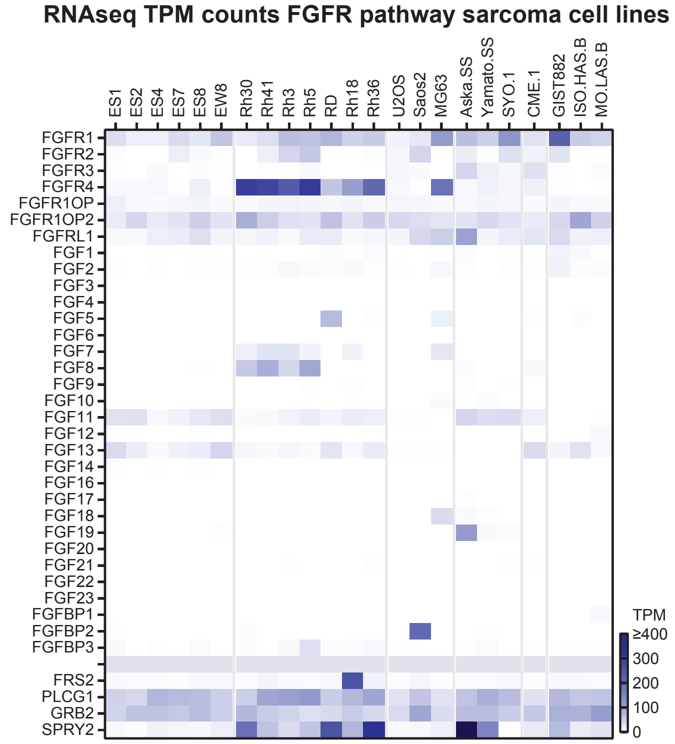


B

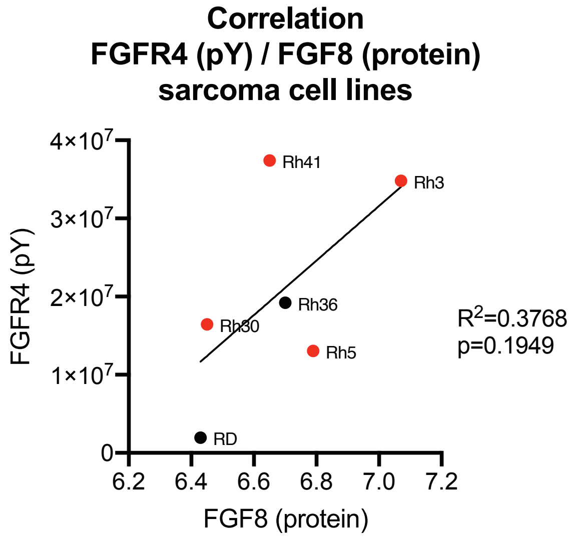

### Supplemental Figure 5

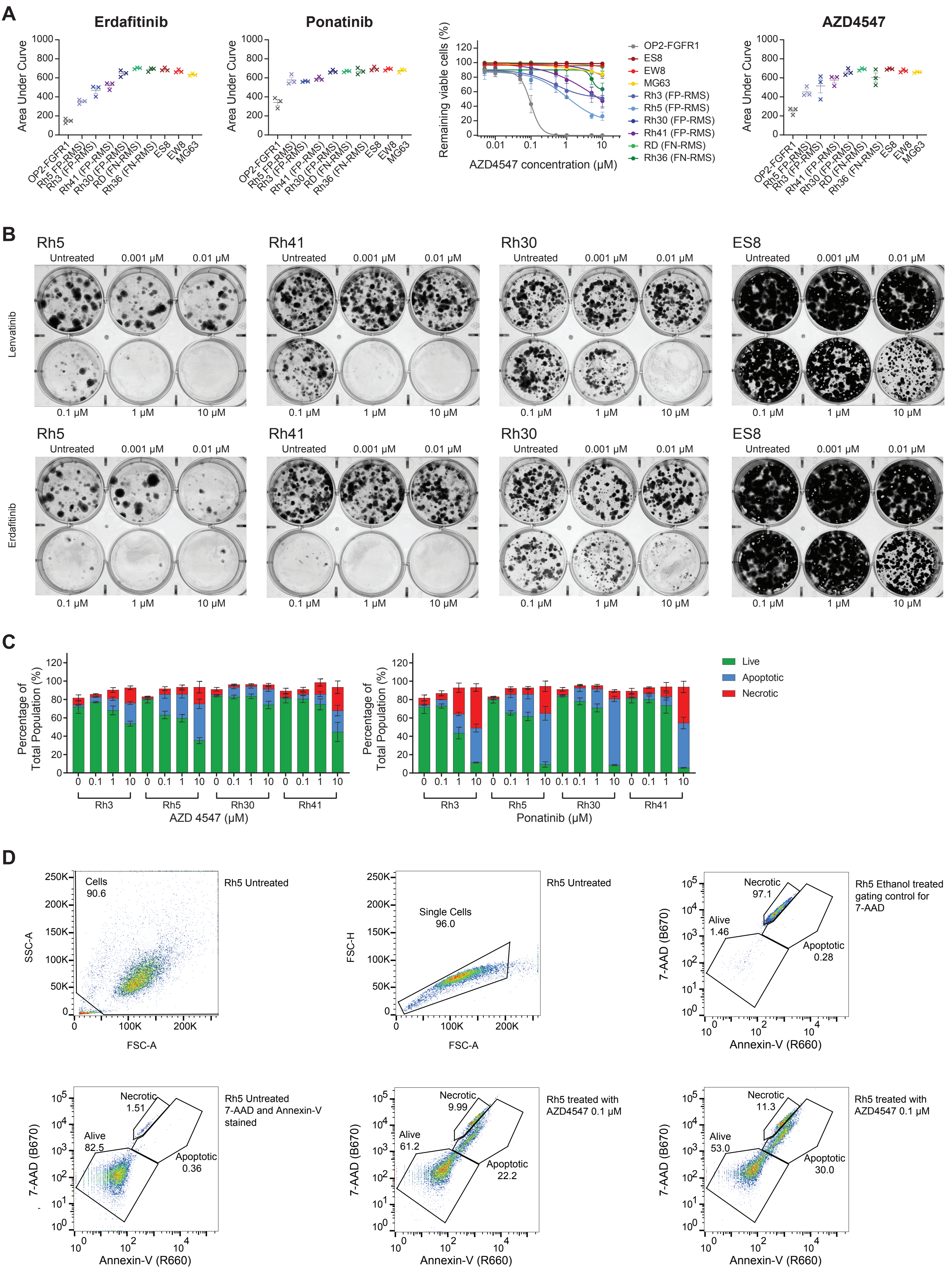

### Supplemental Figure 6

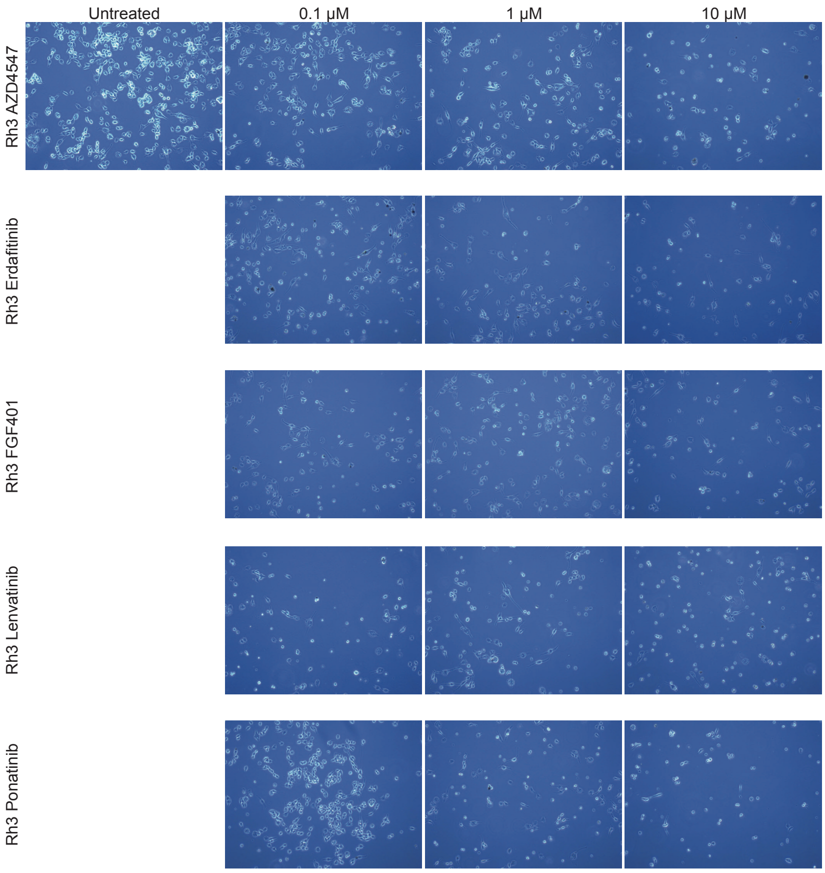

### Supplemental Figure 7

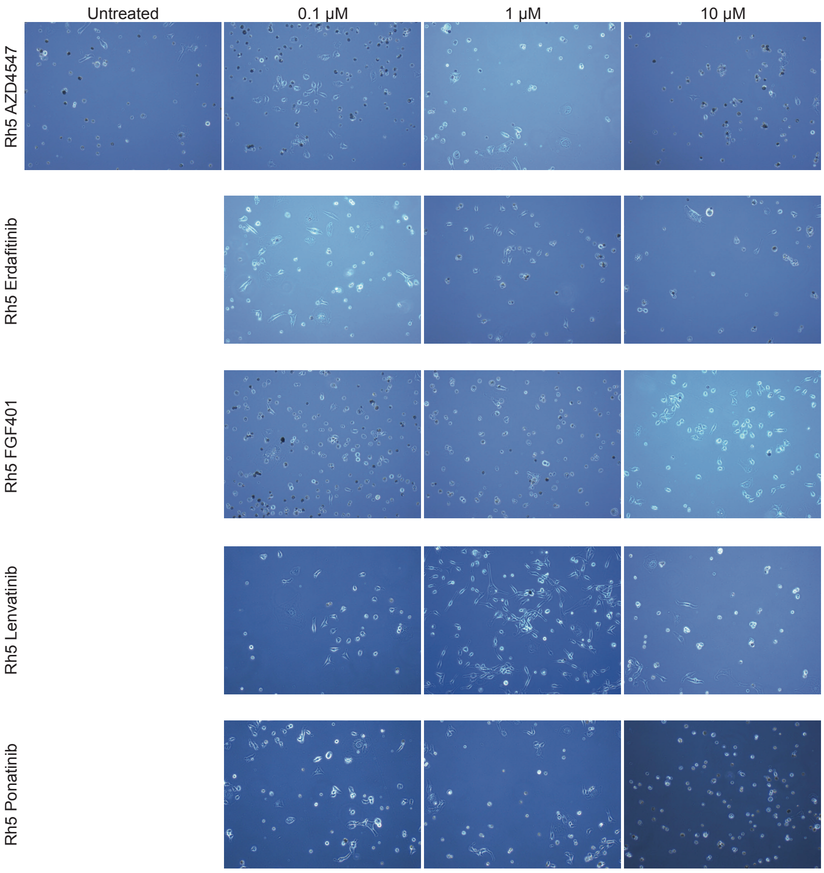

### Supplemental Figure 8

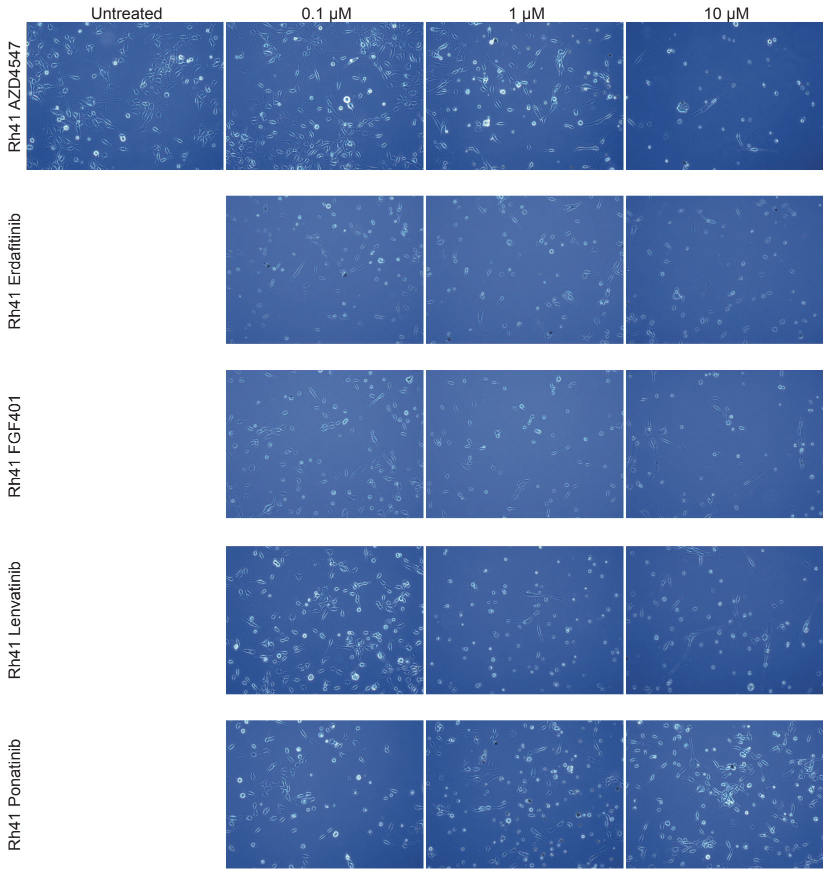

### Supplemental Figure 9

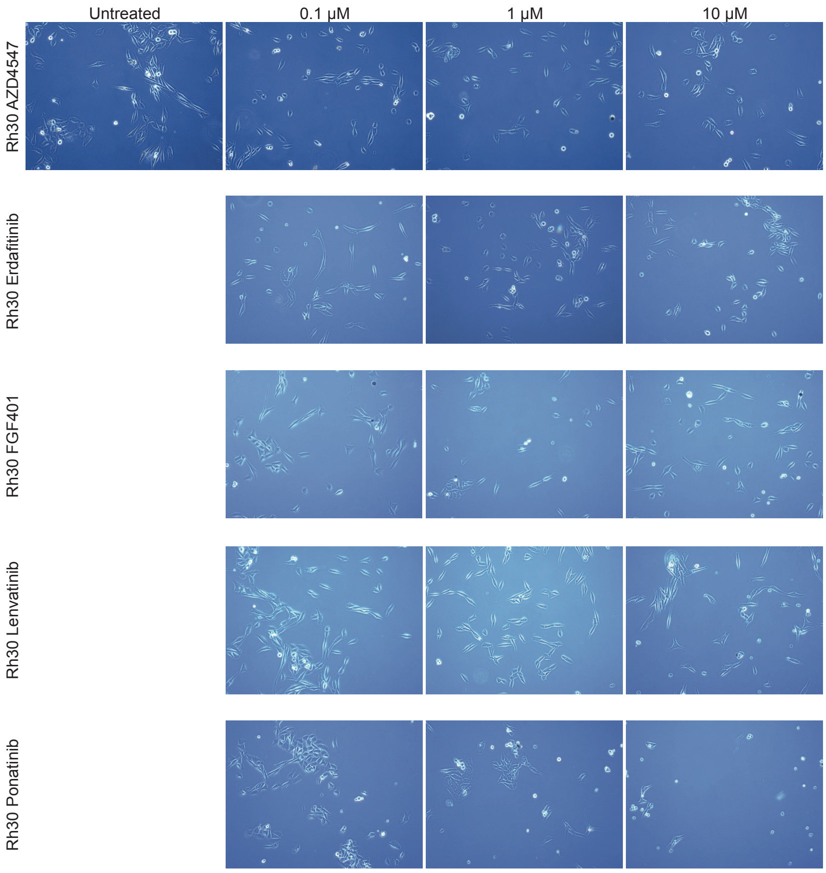

#### Supplemental Figure 10

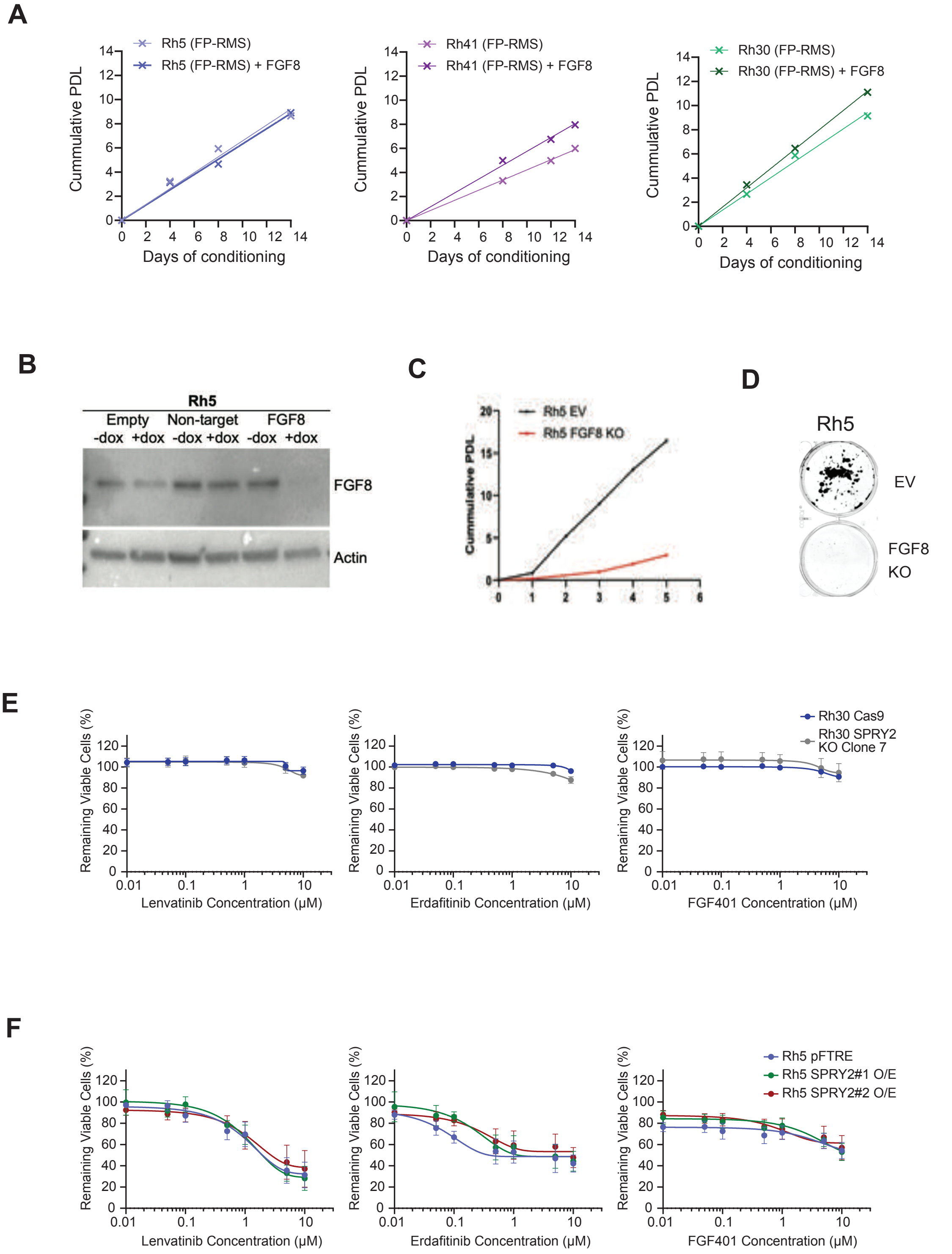

#### Supplemental Figure 11

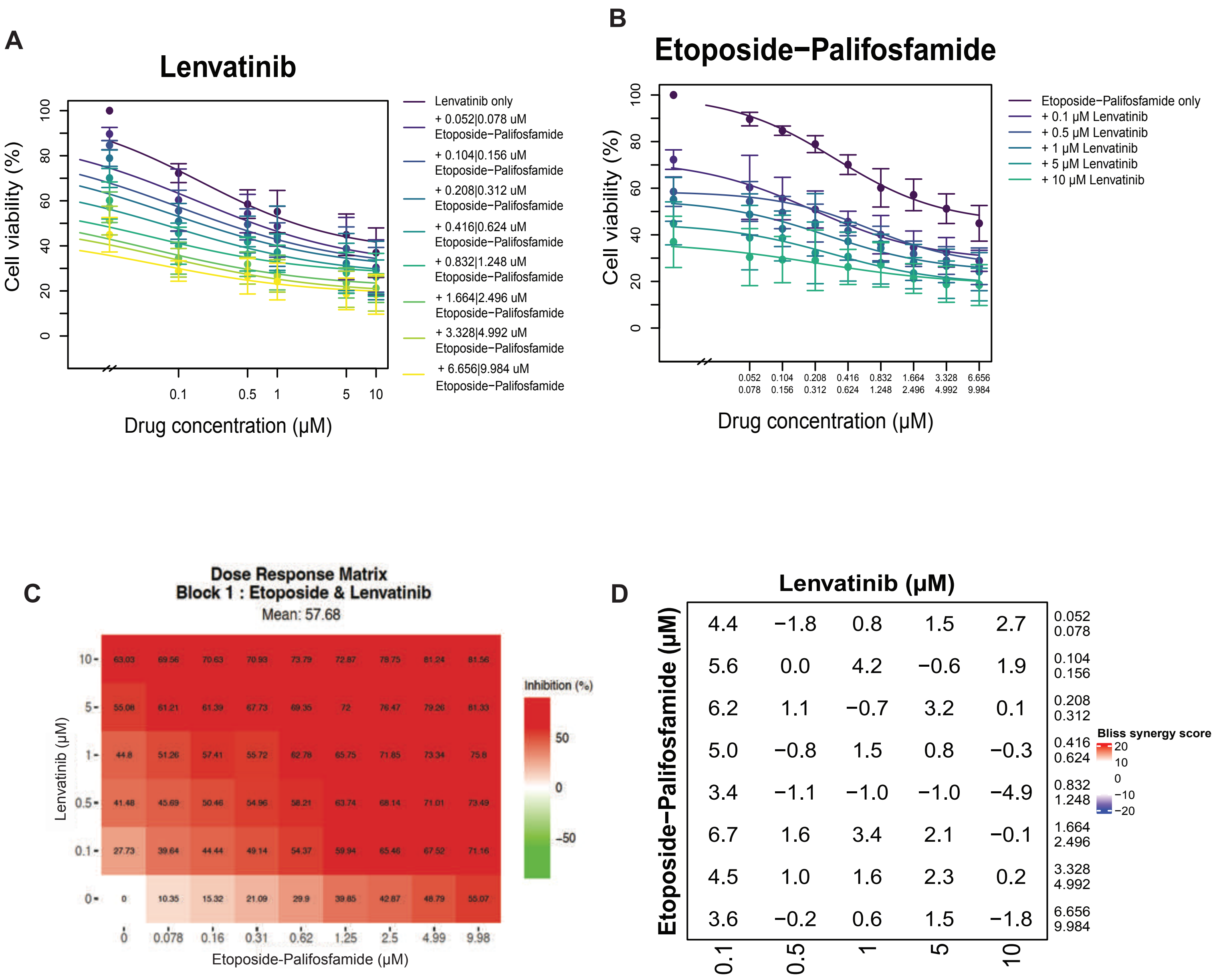

### Supplemental Figure 12

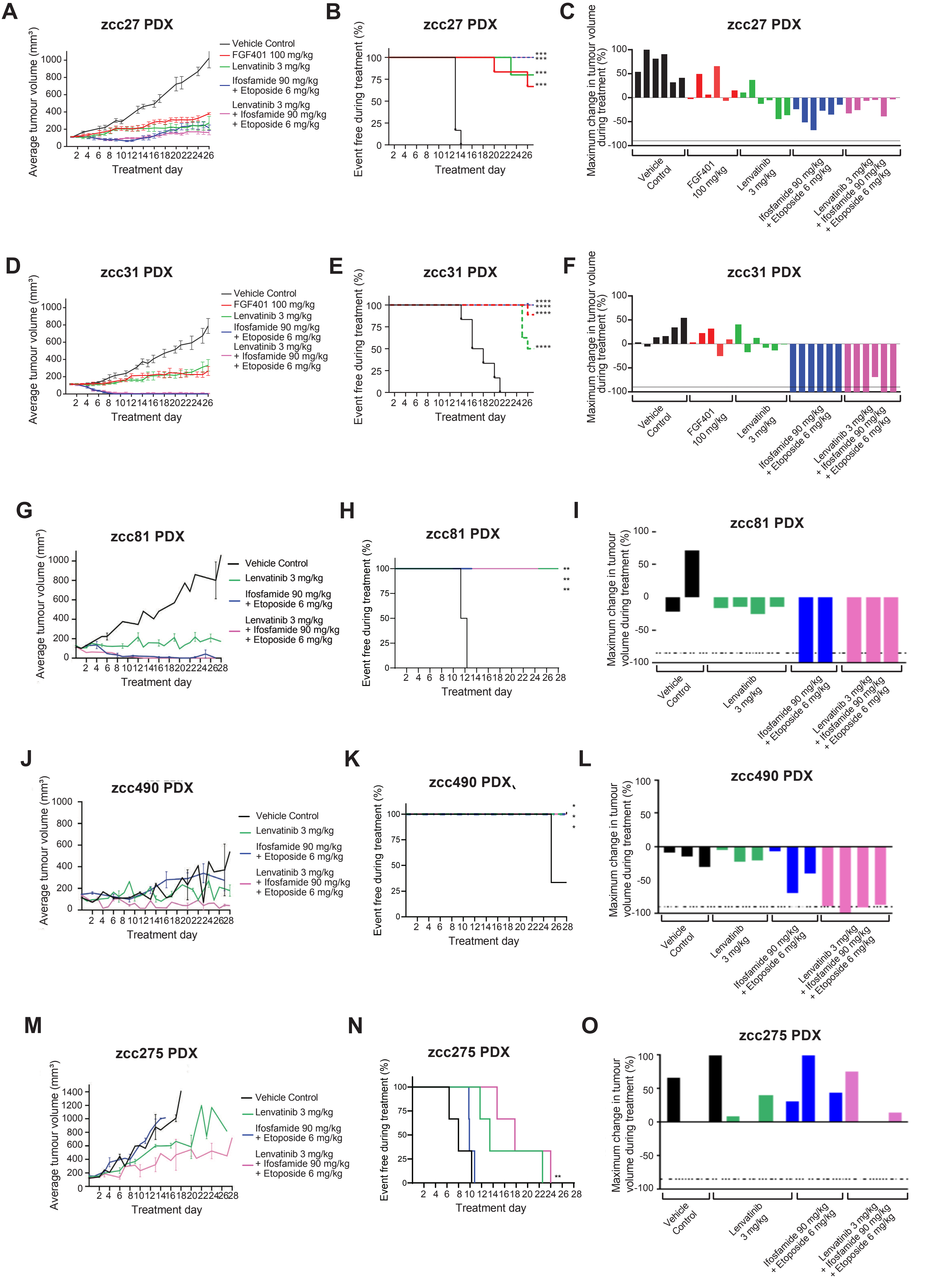
