## Supplemental Table 1 for "Comprehensive multi-platform tyrosine kinase profiling reveals novel actionable FGFR aberrations across pediatric and AYA sarcomas"

| **Cell line** | **Subtype** | **Media and Supplements** | **Culture Type** |
|---|---|---|---|
| OP2-FGFR1 Ba/F3 | (FGFR1 positive control) | RPMI1640, 10% FCS | Suspension |
| ISO.HAS.B | AS | DMEM, 15% FCS | Adherent |
| MO.LAS.B | AS | DMEM, 10% FCS | Adherent |
| ES1 | EWS | RPMI1640, 10% FCS | Adherent |
| ES2 | EWS | RPMI1640, 10% FCS | Adherent |
| ES4 | EWS | RPMI1640, 10% FCS | Adherent |
| ES7 | EWS | RPMI1640, 10% FCS | Adherent |
| ES8 | EWS | RPMI1640, 10% FCS | Adherent |
| EW8 | EWS | RPMI1640, 10% FCS | Adherent. |
| GIST882 | GIST | DMEM, 10% FCS | Adherent |
| MG63 | OS | EMEM, 10% FCS | Adherent |
| Saos2 | OS | Mc Coy's 5A, 10% FCS | Adherent |
| U2OS | OS | DMEM, 10% FCS | Adherent |
| Rh3 | FP-RMS | RPMI1640, 10% FCS | Adherent |
| Rh30 | FP-RMS | RPMI1640, 10% FCS | Adherent |
| Rh41 | FP-RMS | RPMI1640, 10% FCS | Adherent |
| Rh5 | FP-RMS | RPMI1640, 10% FCS | Adherent |
| RD | FN-RMS | DMEM, 10% FCS | Adherent |
| Rh18 | FN-RMS | Mc Coy's 5A, 10% FCS | Adherent |
| Rh36 | FN-RMS | RPMI1640, 10% FCS | Adherent |
| Aska.SS | SynS | DMEM, 10% FCS | Adherent |
| Yamato.SS | SynS | DMEM, 10% FCS | Adherent |
| SYO.1 | SynS | DMEM, 10% FCS | Adherent |
| CME.1 | SynS | RPMI1640, 10% FCS | Adherent |
