## Supplemental Table 2 for "Comprehensive multi-platform tyrosine kinase profiling reveals novel actionable FGFR aberrations across pediatric and AYA sarcomas"

| Patient ID | Final Diagnosis | Disease Stage | Age at Diagnosis | Age at Sample | Vital Status (21 April 2021) | Gender | Somatic SNV (TKs) | Somatic CNV (TKs) | Somatic SV (TKs) | Somatic RNA (TKs) | Recommendation (TKi, from molecular) | Tier |
|---|---|---|---|---|---|---|---|---|---|---|---|---|
| zcc175 | Alveolar soft part sarcoma | Diagnosis | 9 | 10 | Alive | Male | | | | FLT1(1.33); KDR(0.92) | Multi-TKi with VEGFR activity or PAN-VEGFRi for KDR RNA | 1 |
| zcc246 | Alveolar soft part sarcoma | Diagnosis | 21 | 22 | Deceased | Female | | ERBB3(CN:3.21, RNA:-0.05) | | | TARGET sample, no official recommendation | N/A |
| zcc14 | Alveolar soft part sarcoma | Relapse3 | 7 | 17 | Deceased | Female | | | | FLT1(1.47) | VEGFRi inhibitor for VEGFR1 RNA | 1 |
| zcc12 | Angiosarcoma | Diagnosis | 0 | 0 | Alive | Male | | | | FLT1(1.02); FLT4(1.97); KDR(0.92) | VEGFRi for VEGFR1/2/3 RNA | 1 |
| zcc129 | Angiosarcoma | Diagnosis | 6 | 6 | Deceased | Female | NM_002253(KDR): c.2312C>G (p.Thr771Arg) Allele frequency = 0.35 | | | KDR(1.52) | VEGFRi for KDR SNV | 1 |
| zcc87 | Desmoplastic small round blue cell tumour | Diagnosis | 13 | 13 | Deceased | Male | | | | FGFR4(2.53) | Multi-TKi with FGFR activity | 1 |
| zcc364 | Desmoplastic small round blue cell tumour | Diagnosis | 16 | 16 | Alive | Male | | | | | | |
| zcc11 | Ewing's sarcoma | Relapse1 | 14 | 15 | Deceased | Female | | | | | | |
| zcc266 | Ewing's sarcoma | Relapse2 | 13 | 17 | Deceased | Male | | | | JAK1(2.29); KIT(2.09) | No recommendation | N/A |
| zcc20 | Ewing's sarcoma | Relapse2 | 18 | 22 | Deceased | Male | | | | JAK1(4.34) | Non-TKi therapy recommendation or recommendation from other platform | 1 |
| zcc25 | Ewing's sarcoma | Relapse1 | 13 | 17 | Deceased | Female | | | | | | |
| zcc30 | Ewing's sarcoma | Diagnosis | 17 | 18 | Alive | Female | | | | JAK1(4.91) | JAKi for JAK1 RNA | 3 |
| zcc38 | Ewing's sarcoma | Relapse2 | 17 | 20 | Deceased | Male | | | | | | |
| zcc39 | Ewing's sarcoma | Diagnosis | 17 | 17 | Deceased | Male | | | | | | |

| | | | | | | | | | | | | |
|---|---|---|---|---|---|---|---|---|---|---|---|---|
| zcc59 | Ewing's sarcoma | Relapse2 | 8 | 12 | Deceased | Female | | | | | | |
| zcc75 | Ewing's sarcoma | Diagnosis | 8 | 9 | Deceased | Male | | | | | | |
| zcc89 | Ewing's sarcoma | Relapse1 | 12 | 16 | Alive | Female | | ERBB3(CN:3.00) | | | Pan-ERBB inhibitor for ERBB3 CNV | 4 |
| zcc280 | Ewing's sarcoma | Relapse1 | 0 | 2 | Alive | Male | | | | | | |
| zcc121 | Ewing's sarcoma | Relapse1 | 15 | 18 | Deceased | Female | | | | JAK1(3.85) | JAKi for JAK1 RNA | 3 |
| zcc124 | Ewing's sarcoma | Relapse1 | 15 | 17 | Deceased | Male | | | | KIT(2.62) | Non-TKi therapy recommendation or recommendation from other platform | 1 |
| zcc290 | Ewing's sarcoma | Relapse1 | 10 | 11 | Deceased | Male | | | | | | |
| zcc143 | Ewing's sarcoma | Diagnosis | 2 | 2 | Alive | Male | | | | | | |
| zcc148 | Ewing's sarcoma | Relapse2 | 14 | 18 | Deceased | Female | | | | | | |
| zcc326 | Ewing's sarcoma | Diagnosis | 0 | 0 | Alive | Male | | | | | | |
| zcc180 | Ewing's sarcoma | Relapse1 | 17 | 19 | Deceased | Male | | | | | | |
| zcc333 | Ewing's sarcoma | Relapse1 | 9 | 11 | Alive | Male | | | | | | |
| zcc341 | Ewing's sarcoma | Relapse1 | 13 | 14 | Deceased | Female | | FGFR1(CN:7.16) | | | FGFRi for FGFR1 CNV | 2 |
| zcc356 | Ewing's sarcoma | Relapse1 | 12 | 13 | Alive | Male | | | | JAK1(4.20) | JAKi for JAK1 RNA | 3 |
| zcc371 | Ewing's sarcoma | Relapse2 | 4 | 7 | Alive | Female | | | | | | |
| zcc381 | Ewing's sarcoma | Relapse1 | 6 | 8 | Alive | Male | | | | JAK1(3.43) | No recommendation | N/A |
| zcc388 | Ewing's sarcoma | Relapse2 | 3 | 7 | Deceased | Female | | | | JAK1(2.56) | JAKi for JAK1 RNA | 3 |
| zcc389 | Ewing's sarcoma | Diagnosis | 12 | 12 | Alive | Male | | | | | | |
| zcc406 | Ewing's sarcoma | Progression1 | 8 | 9 | Deceased | Male | | | | JAK1(3.62) | No recommendation | N/A |
| zcc210 | Ewing's sarcoma | Diagnosis | 3 | 3 | Deceased | Male | | | | JAK1(2.98) | TARGET sample, no official recommendation | N/A |
| zcc207 | Ewing's sarcoma | Diagnosis | 14 | 14 | Deceased | Female | | | | | | |

| | | | | | | | | | | | | |
|---|---|---|---|---|---|---|---|---|---|---|---|---|
| zcc227 | Ewing's sarcoma | Relapse1 | 0 | 5 | Alive | Male | | | | JAK1(3.75); KIT(2.10) | TARGET sample, no official recommendation | N/A |
| zcc243 | Gastrointestinal neuroectodermal tumour | Diagnosis | 5 | 6 | | Female | | | | | | |
| zcc215 | Gastrointestinal stromal tumour | Progression2 | 10 | 15 | Alive | Male | | | | FGFR1(2.53); KIT(3.82) | FGFRi for FGFR1 RNA, multi-TKi for FGFR1/KIT RNA | 4, 1 |
| zcc419 | Gastrointestinal stromal tumour | Diagnosis | 18 | 18 | Alive | Female | | | | FGFR1(2.18); KIT(2.95) | Multi-TKi with FGFR and KIT activity for FGFR1 RNA and KIT RNA, FGFRi for FGFR1 RNA | 1, 3 |
| zcc275 | High grade undifferentiated sarcoma | Relapse1 | 15 | 15 | Deceased | Male | | | | | | |
| zcc284 | High grade undifferentiated sarcoma | Diagnosis | 2 | 2 | Alive | Male | | | ETV6 - NTRK3 | | NTRKi for ETV6-NTRK3 SV | 1 |
| zcc301 | High grade undifferentiated sarcoma | Diagnosis | 13 | 13 | Deceased | Male | NM_023110.2(FGFR1): c.1638C>A p.(Asn546Lys) Allele frequency = 0.27 | | | | FGFR1i for FGFR1 SNV | 1, 2 |
| zcc359 | High grade undifferentiated sarcoma | Progression2 | 2 | 3 | Deceased | Female | | | | | | |
| zcc418 | High grade undifferentiated sarcoma | Relapse2 | 10 | 17 | Alive | Male | | | | | | |
| zcc211 | High grade undifferentiated sarcoma | Relapse1 | 17 | 17 | Deceased | Male | | | | FLT4(2.19) | TARGET sample, no official recommendation | N/A |
| zcc41 | Infantile fibrosarcoma | Diagnosis | 0 | 1 | Alive | Female | | | ETV6 - NTRK3 | | NTRKi for ETV6-NTRK3 SV | 1 |
| zcc549 | Infantile fibrosarcoma | Relapse1 | 1 | 2 | Alive | Male | | | ETV6 - NTRK3 | | NTRKi for ETV6-NTRK3 SV | 1 |
| zcc209 | Infantile fibrosarcoma | Diagnosis | 0 | 1 | Alive | Female | | | SPECC1L - NTRK3 | | TARGET sample, no official recommendation | N/A |

| | | | | | | | | | | | | |
|---|---|---|---|---|---|---|---|---|---|---|---|---|
| zcc60 | Malignant peripheral nerve sheath tumour | Relapse1 | 13 | 15 | Alive | Male | | | | | | |
| zcc131 | Malignant peripheral nerve sheath tumour | Relapse1 | 15 | 17 | Deceased | Male | | | | | | |
| zcc144 | Malignant peripheral nerve sheath tumour | Diagnosis | 2 | 3 | Alive | Male | | | | | | |
| zcc177 | Malignant peripheral nerve sheath tumour | Diagnosis | 16 | 17 | Deceased | Male | | | | | | |
| zcc187 | Malignant peripheral nerve sheath tumour | Relapse1 | 15 | 16 | Alive | Female | | ALK(CN:9.56, RNA:1.51); EGFR(CN:3.06, RNA:1.26) | | ALK(1.51); EGFR(1.26) | EFGRi for EGFR CNV/RNA | 4 |
| zcc347 | Malignant peripheral nerve sheath tumour | Progression1 | 10 | 12 | Alive | Male | | | | | | |
| zcc28 | Malignant rhabdoid tumour | Diagnosis | 0 | 1 | Deceased | Male | | | | | | |
| zcc36 | Malignant rhabdoid tumour | Diagnosis | 1 | 2 | Deceased | Female | | | | | | |
| zcc113 | Malignant rhabdoid tumour | Diagnosis | 1 | 1 | Deceased | Male | | | | | | |
| zcc161 | Malignant rhabdoid tumour | Diagnosis | 0 | 1 | Alive | Female | | | | | | |
| zcc167 | Malignant rhabdoid tumour | Diagnosis | 0 | 0 | Deceased | Male | | | | c-MET(1.75) | METi for MET RNA | 4 |
| zcc168 | Malignant rhabdoid tumour | Diagnosis | 0 | 1 | Deceased | Female | | | | | | |
| zcc202 | Malignant rhabdoid tumour | Diagnosis | 0 | 1 | Deceased | Male | | | | | | |

| | | | | | | | | | | | | |
|---|---|---|---|---|---|---|---|---|---|---|---|---|
| zcc221 | Malignant rhabdoid tumour | Diagnosis | 3 | 3 | Alive | Female | | | | | | |
| zcc7 | Osteosarcoma | Diagnosis | 16 | 17 | Deceased | Female | | | | | | |
| zcc43 | Osteosarcoma | Diagnosis | 15 | 15 | Deceased | Female | | | | | | |
| zcc97 | Osteosarcoma | Relapse3 | 9 | 15 | Deceased | Female | | | | | | |
| zcc101 | Osteosarcoma | Diagnosis | 13 | 14 | Alive | Female | | | | ROS1(3.03) | Non-TKi therapy recommendation or recommendation from other platform | 3 |
| zcc107 | Osteosarcoma | Progression1 | 16 | 20 | Alive | Male | | | | | | |
| zcc147 | Osteosarcoma | Relapse1 | 12 | 15 | Alive | Male | | | | | | |
| zcc313 | Osteosarcoma | Relapse1 | 15 | 19 | Deceased | Male | | JAK3(CN:4.75) | | | Non-TKi therapy recommendation or recommendation from other platform | 3 |
| zcc322 | Osteosarcoma | Relapse1 | #N/A | 13 | Deceased | Male | | | | | | |
| zcc332 | Osteosarcoma | Relapse2 | 9 | 12 | Alive | Male | | | | | | |
| zcc191 | Osteosarcoma | Relapse1 | 14 | 15 | Deceased | Male | | | | | | |
| zcc351 | Osteosarcoma | Relapse1 | 15 | 17 | Alive | Male | | ERBB4(CN:2.69, RNA:-0.29); ERBB3(CN:3.72, RNA:0.44); ERBB2(CN:3.10, RNA:0.62) | | | pan-ERBBi | No tier |
| zcc386 | Osteosarcoma | Diagnosis | 10 | 10 | Alive | Male | | | LSM1 - FGFR1 | | FGFRi for SV LSM1-FGFR1, Multi-TKi with FGFRi activity for SV LSM1-FGFR1 | 2, 2 |

| | | | | | | | | | | | | |
|---|---|---|---|---|---|---|---|---|---|---|---|---|
| zcc403 | Osteosarcoma | Relapse1 | #N/A | 13 | Alive | Male | | KDR(CN:17.62, RNA:0.69); KIT(CN:16.47, RNA:0.73); PDGFRA(CN:16.61, RNA:1.18) | | JAK2(2.16); PDGFRA(1.18) | TKi with PDGFRA activity for PDGFRA CNV/RNA, JAKi for JAK1 RNA | 1, 3 |
| zcc411 | Osteosarcoma | Diagnosis | 14 | 14 | Alive | Male | | | | | | |
| zcc420 | Osteosarcoma | Diagnosis | 18 | 18 | Deceased | Female | | | | | | |
| zcc470 | Osteosarcoma | Relapse1 | 13 | 13 | Deceased | Female | | | | | | |
| zcc225 | Osteosarcoma | Relapse1 | 15 | 17 | Deceased | Female | | ERBB4(CN:2.89, RNA:0.55) | | ROS1(3.16) | TARGET sample, no official recommendation | N/A |
| zcc410 | Other - Ameloblastic fibrosarcoma | Diagnosis | 18 | 18 | Alive | Male | | | | ABL2(2.78); FGR(1.79); HCK(2.19); LYN(2.36); SRC(2.31) | Dual ABL/SRCi for high ABL2, SRC, LYN, HCK and FGR | 2 |
| zcc19 | Other - Epithelioid sarcoma | Progression3 | 14 | 16 | Deceased | Female | | | | | | |
| zcc330 | Other - Leiomyosarcoma | Diagnosis | 15 | 16 | Alive | Female | | | | | | |
| zcc434 | Other - Low grade fibromyxoid sarcoma | Relapse1 | #N/A | 7 | Alive | Male | | PDGFRB(CN:3.06, RNA:1.43) | | FGFR1(2.26); PDGFRA(0.97); PDGFRB(1.43) | multi-TKI with FGFR and PDGFR activity for RNA/CNV | 1 |
| zcc366 | Other - Round cell sarcoma | Progression1 | 6 | 6 | Alive | Male | | | | | | |
| zcc128 | Other - CIC | Diagnosis | 14 | 15 | Deceased | Female | | | | | | |
| zcc90 | Rhabdomyosarcoma fusion negative | Diagnosis | 7 | 7 | Deceased | Male | | | | FGFR4(1.90) | Non-TKi therapy recommendation or recommendation from other platform | 4 |
| zcc93 | Rhabdomyosarcoma fusion negative | Progression1 | 3 | 5 | Deceased | Male | | | | | | |

| | | | | | | | | | | | | |
|---|---|---|---|---|---|---|---|---|---|---|---|---|
| zcc140 | Rhabdomyosarcoma fusion negative | Relapse1 | 8 | 10 | Alive | Male | | FGFR1(CN:5.01, RNA:2.40) | | FGFR1(2.40) | Non-TKi therapy recommendation or recommendation from other platform | 1 |
| zcc146 | Rhabdomyosarcoma fusion negative | Diagnosis | 4 | 4 | Deceased | Male | | | | | | |
| zcc508 | Rhabdomyosarcoma fusion negative | Relapse1 | 1 | 3 | Alive | Female | | | | FGFR4(2.40) | Non-TKi therapy recommendation or recommendation from other platform | 1, 4, 1, 5, 3 |
| zcc397 | Rhabdomyosarcoma fusion negative | Diagnosis | 15 | 15 | Alive | Female | | | | | | |
| zcc423 | Rhabdomyosarcoma fusion negative | Diagnosis | 15 | 15 | Alive | Male | NM_213647(FGFR4): c.1605C>A (p.Asn535Lys) Allele frequency = 0.43 | | | FGFR4(2.13) | FGFR1i for FGFR4 SNV | 3 |
| zcc136 | Rhabdomyosarcoma fusion negative, spindle cell | Progression1 | 17 | 18 | Deceased | Male | | | | | | |
| zcc23 | Rhabdomyosarcoma fusion positive | Progression1 | 12 | 13 | Deceased | Female | | | | | | |
| zcc27 | Rhabdomyosarcoma fusion positive | Relapse1 | 15 | 18 | Deceased | Male | | | | FGFR4(2.26) | Non-TKi therapy recommendation or recommendation from other platform | 3 |
| zcc31 | Rhabdomyosarcoma fusion positive | Diagnosis | 12 | 12 | Deceased | Female | | | | FGFR4(2.38) | Non-TKi therapy recommendation or recommendation from other platform | 3 |
| zcc40 | Rhabdomyosarcoma fusion positive | Diagnosis | 15 | 16 | Deceased | Male | | | | FGFR4(2.68) | Non-TKi therapy recommendation or recommendation from other platform | 3 |

| | | | | | | | | | | | | |
|---|---|---|---|---|---|---|---|---|---|---|---|---|
| zcc54 | Rhabdomyosarcoma fusion positive | Diagnosis | 9 | 9 | Deceased | Female | | | | FGFR4(2.60) | Non-TKi therapy recommendation or recommendation from other platform | 2 |
| zcc81 | Rhabdomyosarcoma fusion positive | Diagnosis | 16 | 16 | Alive | Female | | | | FGFR4(2.38) | Non-TKi therapy recommendation or recommendation from other platform | 3, 4 |
| zcc102 | Rhabdomyosarcoma fusion positive | Relapse1 | 15 | 17 | Deceased | Female | | ERBB4(CN:2.51, RNA:0.68); ERBB3(CN:2.66, RNA:1.13) | | FGFR4(2.19) | Pan-ErBi for ERBB3 and ERBB4 CNV | 4 |
| zcc118 | Rhabdomyosarcoma fusion positive | Progression1 | 11 | 12 | Deceased | Male | | | | | | |
| zcc153 | Rhabdomyosarcoma fusion positive | Diagnosis | 15 | 15 | Alive | Male | | | | FGFR4(2.45) | Non-TKi therapy recommendation or recommendation from other platform | 1, 4 |
| zcc336 | Rhabdomyosarcoma fusion positive | Diagnosis | 13 | 13 | Deceased | Female | | | | | | |
| zcc360 | Rhabdomyosarcoma fusion positive | Diagnosis | 15 | 15 | Deceased | Male | | | | FGFR4(2.18) | No recommendation | N/A |
| zcc235 | Rhabdomyosarcoma fusion positive | Relapse1 | 17 | 19 | Deceased | Male | | ERBB3(CN:7.98, RNA:1.03); ERBB2(CN:8.07, RNA:-0.49) | | FGFR4(2.57) | TARGET sample, no official recommendation | N/A |
| zcc252 | Rhabdomyosarcoma fusion positive | Relapse2 | #N/A | 17 | Deceased | Female | | | | FGFR4(2.44) | TARGET sample, no official recommendation | N/A |
| zcc53 | Synovial sarcoma | Progression1 | 7 | 8 | Alive | Male | | | | | | |
